# Inflammatory Biomarkers of Asymptomatic and Symptomatic Tuberculosis

**DOI:** 10.1101/2025.10.26.684319

**Authors:** Denis Awany, Dominique T Ariefdien, Simon C Mendelsohn, Virginie Rozot, Humphrey Mulenga, Sarah Nyangu, Michele Tameris, Tumelo Moloantoa, Austin Katona, Fernanda Maruri, Firdows Noor, Ravindre Panchia, Khuthadzo Hlongwane, Kim Stanley, Yuri F van der Heijden, Katie Hadley, Andrew-Fiore Gartland, Craig Innes, William Brumskine, Keertan Dheda, Shameem Jaumdally, Tahlia Perumal, Neil Martinson, Al Leslie, Bernard Fourie, Andriëtte Hiemstra, Stephanus T Malherbe, Gerhard Walzl, Kogieleum Naidoo, Gavin Churchyard, Novel N Chegou, Timothy R Sterling, Mark Hatherill, Thomas J Scriba, the RePORT South Africa Study and the CORTIS Study Teams

**Affiliations:** South African Tuberculosis Vaccine Initiative, Institute of Infectious Disease and Molecular Medicine and Division of Immunology, Department of Pathology, University of Cape Town, Cape Town, South Africa; Perinatal HIV Research Unit, University of the Witwatersrand, Johannesburg, South Africa; Vanderbilt Tuberculosis Center, Vanderbilt University Medical Center, Nashville, TN, USA; South African Medical Research Council Centre for TB Research; Division of Immunology, Department of Biomedical Sciences, Faculty of Medicine and Health Sciences, Stellenbosch University, Cape Town, South Africa; The Aurum Institute, Johannesburg, South Africa; Vaccine and Infectious Disease Division, Fred Hutchinson Cancer Research Center, Seattle, WA, USA; Centre for Lung Infection and Immunity, Division of Pulmonology, Department of Medicine and UCT Lung Institute and South African MRC/UCT Centre for the Study of Antimicrobial Resistance, University of Cape Town, Cape Town, South Africa; Faculty of Infectious and Tropical Diseases, Department of Immunology and Infection, London School of Hygiene and Tropical Medicine, London, UK; Perinatal HIV Research Unit, University of Witwatersrand, Johannesburg, South Africa; Africa Health Research Institute, Durban, South Africa; Division of Infection and Immunity, University College London, UK; Department of Medical Microbiology, University of Pretoria Faculty of Health Sciences, Pretoria, South Africa; Centre for the AIDS Programme of Research in South Africa (CAPRISA), Durban, South Africa; MRC-CAPRISA HIV-TB Pathogenesis and Treatment Research Unit, Doris Duke Medical Research Institute, University of KwaZulu-Natal, Durban, South Africa; School of Public Health, University of the Witwatersrand, Johannesburg, South Africa

**Keywords:** asymptomatic, tuberculosis, transcriptomic, proteomic, biomarker

## Abstract

A large proportion of individuals with tuberculosis (TB) are asymptomatic. The biological and inflammatory underpinnings of asymptomatic TB are unknown and may differ from symptomatic TB. We characterised blood transcriptomic and proteomic profiles in South African community screening vs. health facility-based triage cohorts. Asymptomatic TB shared core transcriptomic and proteomic features with symptomatic TB, including upregulation of innate, interferon and inflammatory pathways and downregulation of T and B cell pathways. Integration of transcriptomic and proteomic data from asymptomatic TB individuals identified two distinct sub-clusters characterized by higher or lower bacterial burden, blood IFN-γ responses, BMI, and chest radiographic abnormalities, suggesting different disease severity. We identified a new blood transcriptomic signature of asymptomatic TB. However, diagnostic performance of transcriptomic and proteomic markers was weaker for asymptomatic TB than symptomatic TB, suggesting that policy development for community-based, asymptomatic TB screening should not adopt biomarkers developed for symptomatic TB triage without further optimization.

## Introduction

More than half of all individuals with tuberculosis (TB) identified in national TB prevalence surveys did not have, recognise, or report typical TB symptoms such, as cough, fever, night sweats, and loss of weight ^1^, and were classified as asymptomatic ^2, 3^. Three studies in South Africa that performed universal community or household contact TB screening, irrespective of symptoms or chest radiographic findings, reported even higher proportions of asymptomatic TB, above 80% ^4, 5, 6^, while a study of clinic attendees in high-risk groups found that 55% of clinic attendees with bacteriologically-confirmed TB were asymptomatic^7^. Recent studies also suggest that symptoms, such as cough, are not required for aerosol release of *M. tuberculosis* ^8, 9^, raising the possibility that asymptomatic TB may contribute to transmission. Indeed, modelled projections based on data from 25 historic cohorts suggest that 52% of *M. tuberculosis* transmission was from individuals with asymptomatic disease ^10, 11^. Thus, asymptomatic TB is not only common, but finding, treating and preventing asymptomatic may be important for the success or failure of global TB control measures.

Yet, a number of fundamental questions remain unanswered, including whether asymptomatic TB differs from symptomatic disease in its biology, severity, morbidity, risk of post-TB lung disease or mortality, and likelihood of detection by host biomarkers and molecular diagnostic tests. A critical unknown for global health policy-makers is how diagnostic tools originally developed for health facility-based triage of persons presenting with symptoms suggestive of TB to primary health clinics will perform for community-based screening of asymptomatic TB. A major obstacle is that those with asymptomatic TB do not self-present for care and are not detected by active symptom screening in facilities or communities. As a result, asymptomatic TB is poorly understood and the host immune and inflammatory responses in asymptomatic TB remains to be characterized. We recently assessed several published blood transcriptomic signatures of TB with promise as tests for TB triage tests ^12^, predicting progression to TB disease ^13, 14, 15, 16^, and diagnosis, in the CORTIS studies. We observed promising diagnostic performance among symptomatic participants for many signatures, but specificity for asymptomatic TB was sub-optimal. At 90% sensitivity, the specificity for all signatures fell below 50% ^17^, which falls well short of the minimum criteria set out in the Target Product Profile (TPP) for a TB screening test ^18^. New symptom-agnostic tools are needed for community based TB screening. However, if asymptomatic TB in the community differs in underlying mechanism and/or severity from symptomatic TB detected at health facilities, different screening approaches may be needed.

Radiological imaging in close contacts of TB cases suggest a large range of pulmonary and other thoracic manifestations and ^18^F-Fluorodeoxyglucose positron emission and computed tomography (PET/CT) imaging identifies a high degree of variability in inflammatory and metabolic activity ^19^. Importantly, individuals with PET-CT active lesions were at significantly higher risk of subsequent progression to bacteriologically-positive TB than those with PET-CT inactive lesions ^19^. Studies that screen for asymptomatic TB among contacts thus may include individuals with early disease, who are sampled while progressing from *M. tuberculosis* infection to symptomatic TB, while others have undulating disease that cycles through asymptomatic and symptomatic stages^20^. Like symptomatic TB^21^, asymptomatic TB may encompass a spectrum of disease states with distinct biological and inflammatory responses.

Here, we characterised the inflammatory response in individuals with bacteriologically-confirmed, asymptomatic TB using blood transcriptomic and proteomic profiling, and compared these profiles to symptomatic TB in contemporaneous cohorts. We also sought to identify a blood transcriptomic signature of asymptomatic TB. We show that asymptomatic TB was characterised by increases in inflammatory responses and innate immune activation previously described for symptomatic TB. However, these perturbations were less pronounced in asymptomatic TB than symptomatic TB, which may have profound implications for the success of new diagnostic tools. In addition, asymptomatic TB was a highly heterogenous, intermediate phenotype in the TB spectrum comprised of at least two sub-clusters, rather than a distinct phenotype. The findings suggest that host biomarkers and molecular tests developed and validated for facility-based triage of symptomatic TB cannot simply be re-purposed for community-based screening of asymptomatic TB without further optimization.

## Methods

### Study design and participants

#### Regional Prospective Observational Research for Tuberculosis (RePORT) South Africa Cohorts

The Regional Prospective Observational Research for Tuberculosis (RePORT) South Africa (SA) network enrolled two parallel cohorts to study novel diagnostic biomarkers, using a single standardized protocol, at six clinical sites in districts with high TB incidence rates. The first, community-based screening cohort (**Figure 1A**), which includes adult (≥18 years) household contacts (HHC) of index cases with bacteriologically-confirmed pulmonary TB (HHC Screening Cohort), has been described^6^. The second, facility-based triage cohort (**Figure 1B**) includes patients presenting at healthcare facilities with presumptive TB, all of whom were symptomatic (Clinic Triage Cohort).

**Figure 1.**
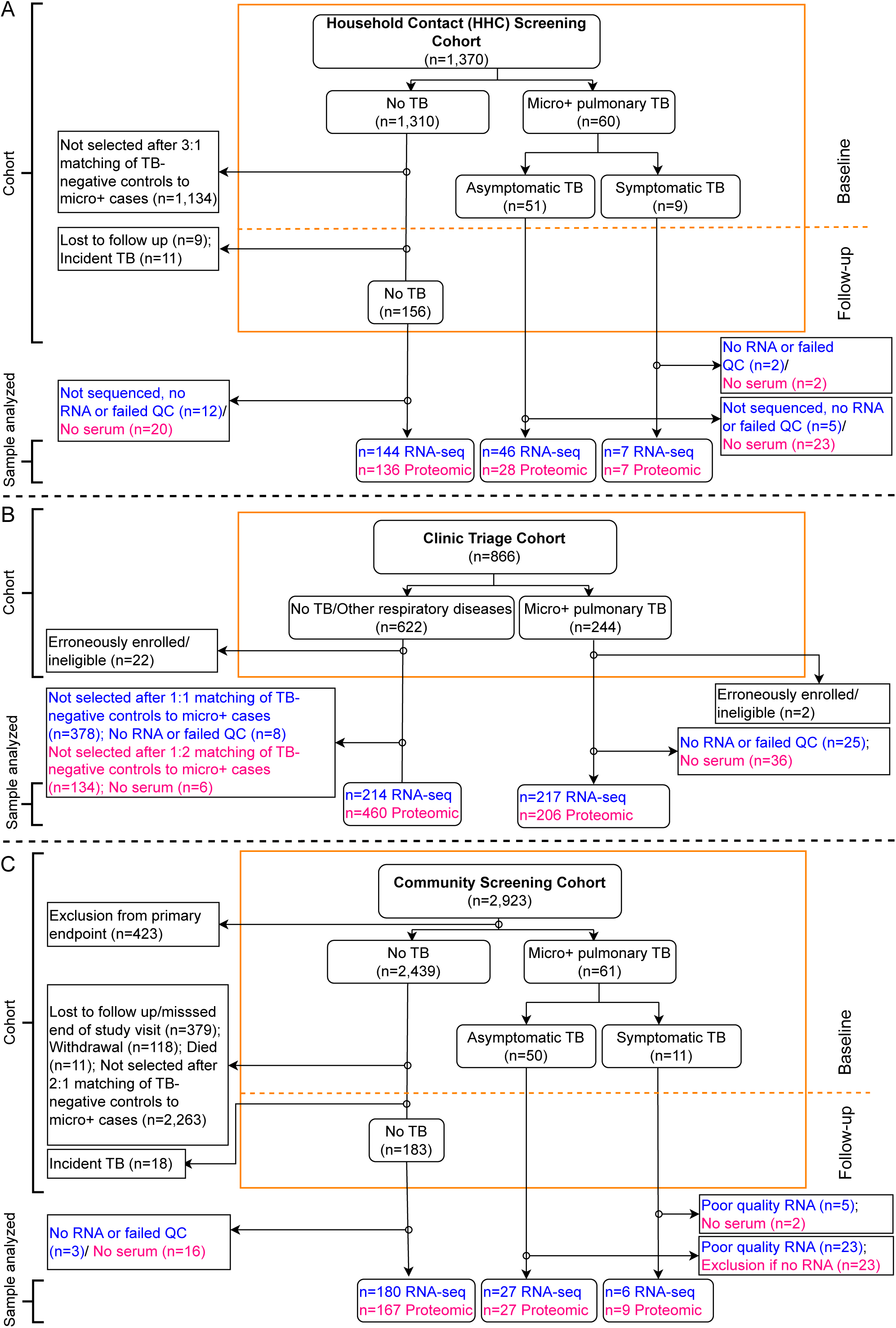
Participant and sample selection for the transcriptomic and proteomic analyses in the three cohorts. The diagrams illustrate enrolment, classification of TB cases and controls and inclusion and exclusion of samples for transcriptomic (blue) or proteomic (red) analyses for the (A) Household Contact (HHC) Screening Cohort, (B) Clinic Triage Cohort, and (C) Community Screening Cohort. For each cohort, stratification into asymptomatic TB, symptomatic TB, and TB-negative (no TB) is shown. Subsequent exclusions for reasons such as loss to follow-up, poor sample quality (“failed QC”), and case-control selection are indicated, leading the final number of samples analysed for RNA-seq and proteomics for each group.

For all participants, symptom screening, chest radiography (CXR), and spontaneously- expectorated sputum for Xpert MTB/RIF Ultra (Cepheid, CA, USA), liquid culture (Mycobacteria Growth Indicator Tube [MGIT], BACTEC, Beckton Dickinson, NJ, USA), and smear microscopy, were performed at baseline, regardless of symptom or CXR status. Blood was collected in PAXgene Blood RNA tubes (PreAnalytiX, Hombrechtikon, Switzerland) for RNA analysis and BD Vacutainer serum-separating tubes (Franklin Lakes, NJ, United States) for serum analysis. A participant was classified symptomatic if they reported one or more of cough, fever, weight loss, fatigue, night sweats, pleuritic chest pain, or haemoptysis, for any duration. CXR were read once by an experienced investigator at each site using a standardised form and recorded as normal or abnormal (compatible with pulmonary TB), based on absence or presence of any cavitation, opacity, mediastinal or hilar adenopathy, pleural effusion, bronchiectasis, or collapsed lung.

Bacteriologically-confirmed pulmonary TB disease was defined as one or more sputum specimens positive by MGIT culture or Xpert Ultra (excluding trace positive). Sputum induction was not performed. Participants who were unproductive of sputum, or with Xpert Ultra trace positive result only, were considered sputum-negative in the primary analysis. Participants who, after investigation for TB at baseline, were not confirmed by positive sputum MGIT culture or Xpert Ultra, were defined as controls for the purpose of diagnostic analyses. Prevalent TB was defined as cases diagnosed on sputum samples collected at the baseline visit. Symptomatic and asymptomatic TB were defined as bacteriologically-confirmed prevalent TB with or without reported symptoms, respectively. All participants diagnosed with TB disease were referred for treatment.

### Community Screening Cohort (CORTIS)

The CORTIS study has been described^4^. Briefly, we performed community-based, universal sputum screening of HIV-uninfected adults residing in TB-endemic communities in South Africa, as for RePORT SA (**Figure 1C**). The study enrolled 2,923 participants, who were pre-screened to enrich the study population for individuals who tested positive for the RISK11 transcriptomic signature. Prevalent TB diagnosed at baseline was microbiologically-confirmed in at least 2 sputum samples.

### Case-control design

Biomarker analyses were performed on case-control subsets. For all cohorts, all TB cases were included from all cohorts (**Figure 1**). For the HHC Screening Cohort, two controls were selected for each TB case by propensity score matching based on site, sex, age, and HIV status (**Figure 1A**). For the Clinic Triage Cohort, one and two controls were selected for each TB case for respectively the RNA-seq and proteomic analyses, respectively, based on site, sex, age, BMI and HIV status (**Figure 1B**). For the Community Screening Cohort three controls were selected for each prevalent TB case by propensity score matching based on site, sex and age (**Figure 1C**). The different ratios of cases to controls were informed primarily on study design, but operational feasibility and available funding at the time of the analyses was also considered.

### RNA-sequencing

For RePORT SA, participant RNA was extracted manually from PAXgene tubes using the PAXgene blood RNA kit (PreAnalytiX, Hombrechtikon, Switzerland). For CORTIS participants, RNA was extracted using a high-throughput, standardised, fully automated procedure on the Freedom EVO 150 robotic platform (Tecan, Männedorf, Switzerland) with the Maxwell SimplyRNA kit (Promega, Madison, WI, USA). RNA was treated using the NEBNext Globin & rRNA Depletion Kit (New England Biolabs, Ipswich, MA, United States) and sequenced on the DNBEQ-G400 sequencing platform, yielding at least 30 million paired-end 100-base reads per sample.

### Blood transcriptome analyses

#### RNA-seq Data Processing and Differential Expression Analysis

Raw sequencing reads were quality-checked using FastQC^22^ and MultiQC^23^. Trim-RNA Galore was used to remove adapters and filter raw reads below 36 bases long, and leading and trailing bases below quality 20. The filtered reads were aligned to the *Homo sapiens* reference genome (GRCh38.p14) using the HISAT2 v2.2.1 aligner ^24^. The aligned reads were quantified to obtain gene-level counts using featureCounts from Subread v2.1.0^25^. Lowly expressed genes with counts per million (CPM) > 2 in at least 5 samples were retained. Differentially expressed genes (DEGs) were identified using DESeq2, applying a false discovery rate (FDR) threshold of <0.05 and an absolute *log*_2_ fold change >1. For subsequent analyses requiring normalized expression, voom transformation was applied to the raw count data to yield *log*_2_-normalized expression data.

### Molecular Distance to Health

We calculated molecular distance to health (MDH) scores, which quantified the deviation from a sample’s gene expression profile from a healthy reference population, as previously described^26^. In summary, we filtered the normalized gene expression and retained the 75% most variable genes, and the calculated median and standard deviation of the gene expression for the healthy control samples. Next, a z-score normalization was performed using the control samples as reference. Absolute values of the normalized z-scores were obtained and values above a threshold of 2 was set to zero. Finally, MDH scores were calculated as the average of these absolute z-scores for the perturbed genes in a sample.

### Pathway enrichment

To identify differentially regulated pathways, we performed differential pathway analysis using quantitative set analysis of gene expression (QuSAGE^27^) with Blood Transcription Modules (BTM) gene sets representing diverse biological contexts. To complement the pathway analysis, we also employed gene co-expression network analysis using CEMiTool^28^ to identify sets of coherently expressed gene modules and performed over-representation analysis (ORA) for genes within each module using the clusterProfiler R package^29^. Pathways with FDR-adjusted p-values <0.05 were considered significantly enriched.

### Measurement of serum proteins

Serum proteins were measured using Bio-Plex multiplex bead arrays on the Bio-Plex platform (Bio-Rad Laboratories, CA, USA), as previously described^30^. Samples from the RePORT SA HHC Screening and Clinic Triage cohorts were assayed using the Bio-Plex Pro Human Cytokine 27-Plex Panel and the Bio-Plex Pro Human Apolipoprotein 10-plex Assay Panel. Samples from the CORTIS Community Screening cohort were assayed using the Bio-Plex Pro Human Cytokine, Chemokine, and Growth Factor 48-Plex Panel.

### Machine Learning for Biomarker Discovery

To identify a novel host transcriptomic signature of asymptomatic TB, we applied an in-house machine learning pipeline—OmicScan (https://github.com/SATVILab/OmicScan)—to analyse the combined datasets from the HHC Screening and Community Screening cohorts. We split the combined dataset into a signature discovery set (75%) and a hold-out validation set (25%), and then further partitioned the discovery set into training (20%) and test (20%) sets (see Supplementary Methods for details).

The OmicScan pipeline included gene prefiltering to remove noisy/uninformative genes, gene ranking to retain the top outcome-associated genes, and derivation of the parsimonious biomarker model. Next, we applied four machine learning (ML) algorithms—Support Vector Machine (SVM), Random Forest (RF), Gradient Boosting (XGBoost) and K-Nearest Neighbour (KNN)—to train predictive models and subsequently evaluated their performance on the discovery test set. We used a “g-score”, a weighted combination of SHAP (Shapley Additive exPlanations) value and permutation importance score, to rank genes from each ML model based on their contribution to the predictive model. Together, the g-score determined a weighted usefulness of the gene.

Finally, we performed a greedy forward search on the top 10 ranked genes to derive the most predictive, parsimonious combination of genes for diagnostic application (Supplementary Methods).

### Integration of transcriptomic and proteomic data to discover clusters

We sought to integrate the abundance of serum proteins and blood RNA-seq datasets from asymptomatic TB cases from the two Screening cohorts to understand the heterogeneity amongst this TB phenotype. We employed Similarity Network Fusion (SNF) in SNFTool^31^ to integrate transcriptomic and proteomic data modalities, allowing unsupervised identification of molecular subgroups within the asymptomatic TB dataset. Optimal cluster numbers were identified (2 to 8 clusters) by eigen-gap and rotation cost methods^31^. After identifying subclusters, we utilized the *clValid* R package^32^ to assess cluster stability and validity through several metrics, including the average proportion of non-overlap (APN), the average distance between means (ADM), and the adjusted Rand Index (ARI) (Supplementary Methods).

To explore the clinical relevance of the subclusters identified, subcluster membership was tested for association with TB-related clinical features, including liquid culture time to positivity (TPP), Xpert Ultra classification, sputum smear microscopy grade, CXR abnormality score generated by Qure.ai (Bengaluru, Karnataka, India), body-mass index (BMI), and IFN-γ levels obtained for TBAg1 and TBAg2 tubes in the QuantiFERON-TB Plus (QIAGEN, Hilden, Germany) IGRA. Group comparisons were made using Wilcoxon rank-sum tests for continuous variables and chi-squared tests for categorical variables.

### Ethical approval

The RePORT SA and CORTIS study protocols were approved by the institutional research ethics committees at each participating site; the RePORT SA protocol was also approved at Vanderbilt University Medical Center, USA. All participants provided written, informed consent prior to participation.

## Results

### Participant characteristics

Among HHC Screening Cohort participants, RNA-seq data for blood transcriptomic profiling was available for 46 asymptomatic TB cases, 7 symptomatic TB cases, and 144 controls (**Figure 1A**). Among Clinic Triage Cohort participants, RNA-seq data was available for 217 symptomatic care-seeking TB cases and 214 controls without TB, who likely had other respiratory diseases (ORD, **Figure 1B**). Among Community Screening Cohort participants, RNA-seq data was available for 27 asymptomatic TB cases, 6 symptomatic TB cases, and 180 controls (**Figure 1C**). The demographic characteristics of the participants are summarized in **Supplementary Figure** and **1 Supplementary Table 1**.

### Transcriptomic profiles of asymptomatic TB

We first computed the Molecular Distance to Health (MDH)^33^, a composite score of the number of transcripts in the transcriptome that significantly differ from healthy controls and the degree of the difference in expression levels. Symptomatic TB cases who presented to health facilities (clinic triage) had markedly higher MDH scores than those without TB, who had ORD (**Figure 2A**). Strikingly, although individuals with asymptomatic TB in both Screening cohorts had significantly higher MDH scores than TB-negative controls, these scores were much lower than in symptomatic TB in the Clinic Triage Cohort (**Figure 2A**). This suggests that asymptomatic TB may be characterised by less perturbation of blood leukocyte gene expression and lower inflammatory responses and suggests that those with asymptomatic TB may have less severe disease. The absolute number of differentially expressed genes also supported this pattern (**Supplementary Figure 2**). To assess this hypothesis further, we compared microbiological features between these groups. Symptomatic TB cases among the Clinic Triage Cohort also had lower BMI, and lower time to culture positivity or were more likely to have higher Xpert Ultra semiquantitative results (both signifying higher bacillary load), than asymptomatic TB cases (**Supplementary Figure 2**), further supporting that those with asymptomatic TB may have less severe disease.

**Figure 2.**
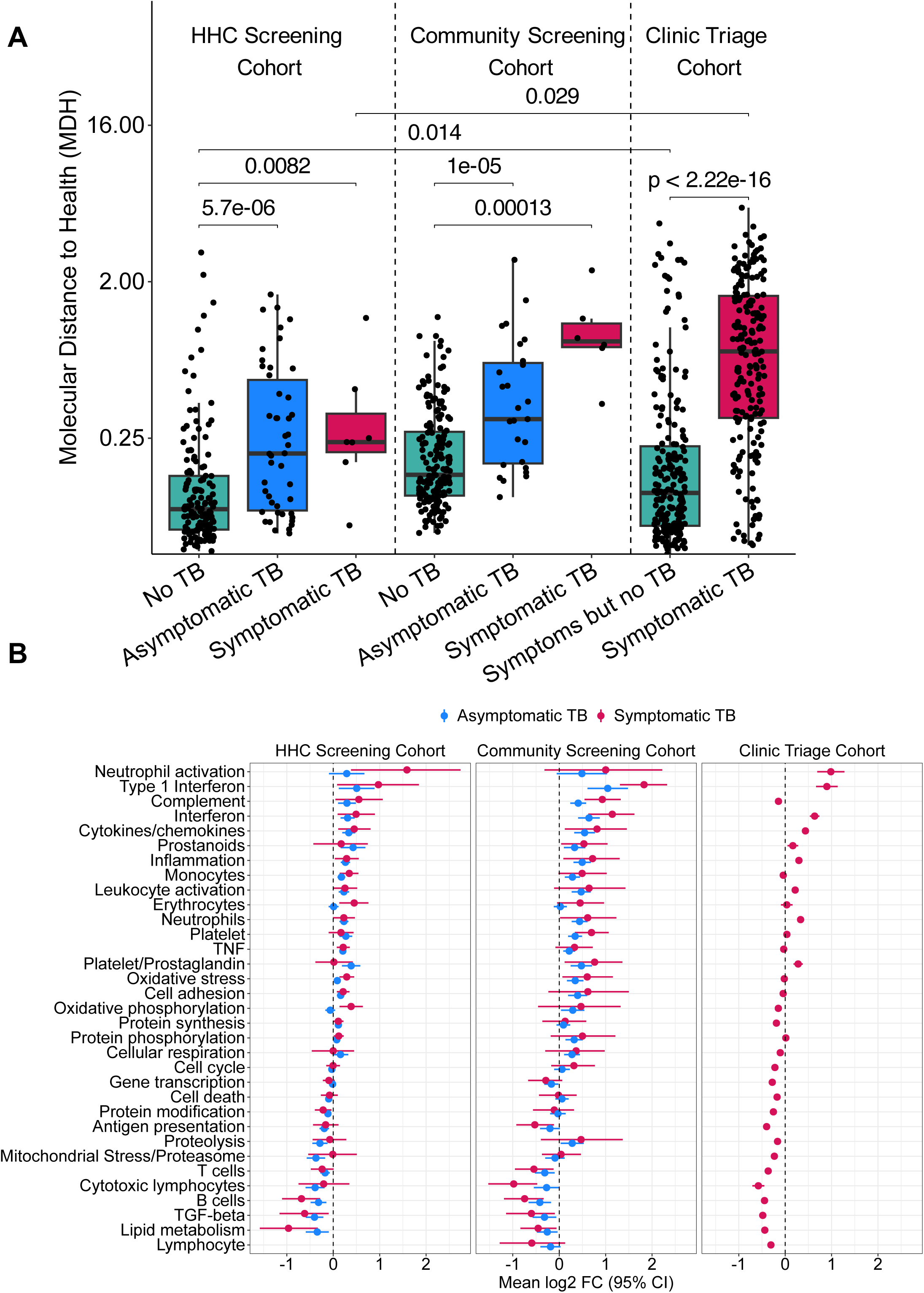
Blood transcriptome perturbation and pathways, biological processes, and gene sets modulated in symptomatic and asymptomatic TB. (**A**) Box and whisker plots showing molecular distance to health, a composite score that quantifies transcriptome-wide deviation from healthy controls, integrating both the number and magnitude of significantly regulated gene transcripts. On the left are Household Contact (HHC) Screening participants without TB, with asymptomatic TB or symptomatic TB. In the middle are Community Screening participants without TB, with asymptomatic TB or symptomatic TB. On the right are Clinic Triage participants without TB or bacteriologically-confirmed symptomatic TB. The horizontal lines indicate the median, the boxes the interquartile range (IQR) and the whiskers the range of data with 1.5 x *IQR* from the lower and upper quartiles. The shown p-value was computed with the Mann-Whitney U test. (**B**) Differential gene set enrichment analysis using QuSAGE on blood RNA-sequencing data, for the gene sets/modules listed on the left, from individuals with asymptomatic TB or symptomatic TB, relative to TB-negative controls, in the HHC Screening Cohort (left), Community Screening Cohort (middle), and Clinic Triage Cohort. Dots denote the enrichment score estimate and error bars the 95% confidence interval. The order of the gene sets is based on ranking from highest to lowest enrichment score estimates in prevalent TB (symptomatic and asymptomatic combined) from the HHC Screening Cohort.

To explore pathways and processes of the transcriptional host response during asymptomatic TB, we performed gene set enrichment analysis (GSEA) using blood transcriptional modules (BTM)^34^ contrasting asymptomatic TB transcriptomes with those from TB-negative controls (**Figure 2B**) among the HHC Screening and Community Screening cohorts. In both these cohorts, asymptomatic TB was associated with significantly elevated gene modules that reflect innate, inflammatory, and myeloid processes, including interferon (IFN) and type I IFN, cytokines and chemokines, inflammation, complement activation, neutrophil activation, leukocyte activation, monocytes, prostanoids, platelet/prostaglandin, and TNF. Cellular processes and metabolism pathways, such as oxidative stress, protein phosphorylation, oxidative phosphorylation, and cellular respiration, were also elevated in asymptomatic TB, relative to TB-negative controls. By contrast, lymphoid pathways, including B cells, T cells, cytotoxic T lymphocytes, lipid metabolism, and TGFβ pathways were significantly lower in asymptomatic TB (**Figure 2B)**. Broadly similar patterns where observed when symptomatic TB detected among Clinic Triage participants was contrasted with TB-negative ORD controls, as extensively reported when symptomatic TB has been compared to TB-negative controls^33, 35, 36, 37^. However, it was notable that complement activation, monocytes, platelets, TNF, and cellular processes and metabolism pathways were not significantly elevated. This is likely because TB cases were compared to symptomatic, care-seeking controls with other respiratory diseases, rather than healthy controls, which have been a more commonly used comparator group for gene module and pathway analyses in published studies.

In light of the consistent patterns observed in the Community HHC Screening and Community Screening cohorts, and to provide a larger sample size to explore cellular and inflammatory processes across the TB disease spectrum, we combined RNA-seq data from these cohorts and performed weighted gene co-expression network analysis (WGCNA), a clustering method that reduces high dimensional data, such as full transcriptomes, into modules that preserve intrinsic relationships between variables within a network structure ^28, 38^.

WGCNA using the top 75% most variable genes identified six modules for which enrichment scores could be computed (**Figure 3A**). When aligned along the TB spectrum, spanning no-TB controls, asymptomatic TB, and symptomatic TB, which were further stratified into Community Screening and Clinic Triage cohorts, enrichment scores for M2, M3, and M4 increased progressively along this trajectory, while enrichment scores for M5 and M6 decreased progressively (**Figure 3A**).

**Figure 3.**
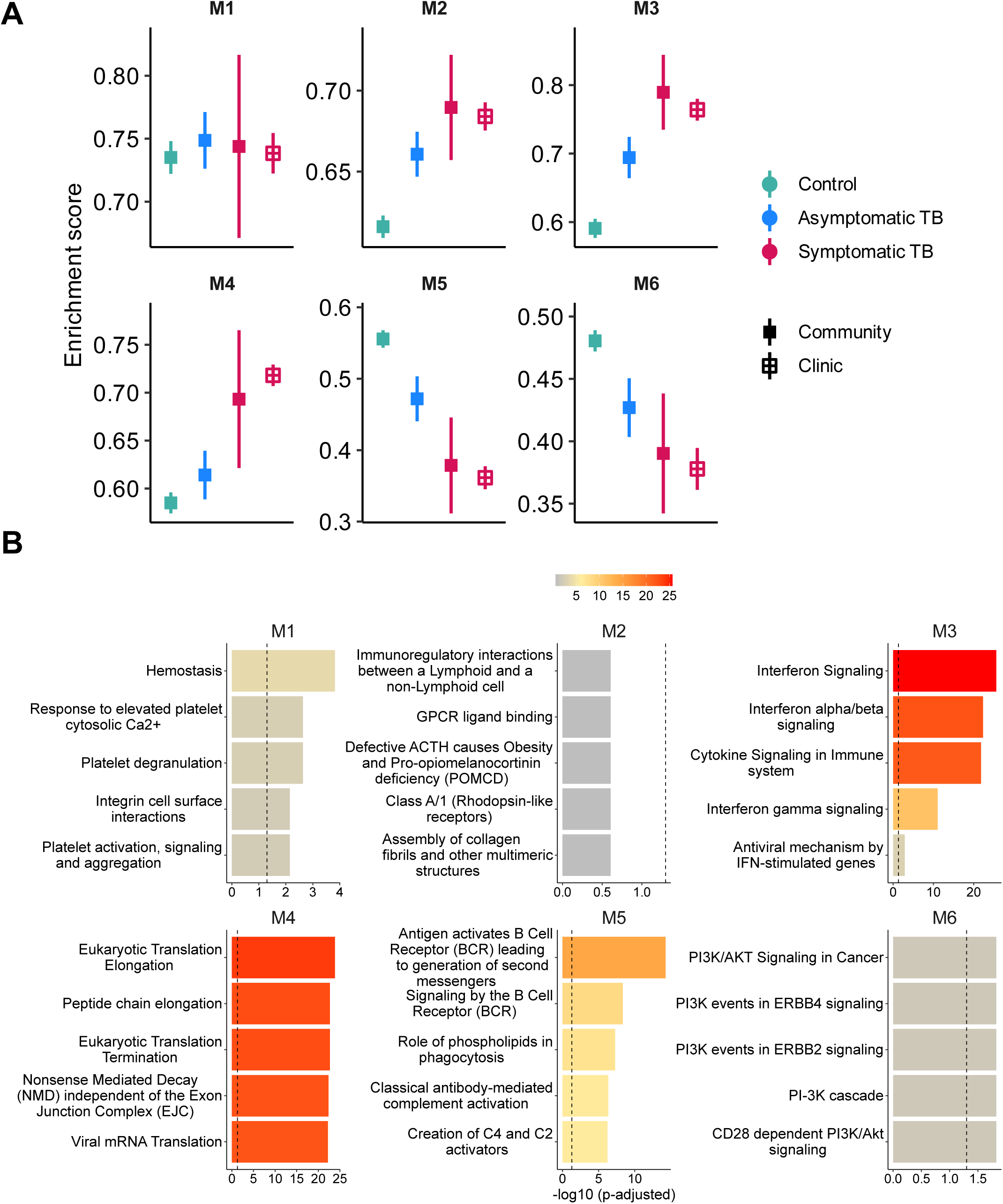
Blood transcriptome perturbations increase progressively along the TB spectrum. Analysis of RNA-seq data from individuals categorised and ranked along the TB spectrum, from healthy controls, asymptomatic TB, symptomatic TB in Community Screening participants and symptomatic TB in Clinic Triage participants, by weighted gene co-expression network analysis using CEMiTool. For these analyses, data from the HHC Screening (RePORT SA) and Community Screening (CORTIS) cohorts were combined. (**A**) M1 through M6 represent six modules of co-expressed gene networks identified by CEMiTool. Dots denote the mean enrichment score estimate and error bars the 95% confidence interval. (**B**) Gene set overrepresentation analysis for the identified gene modules. Displayed are the top five significantly enriched pathways and processes for each of the six modules, ranked within each module according to Benjamini-Hochberg-adjusted *p*-value. The colour intensity for each module corresponds to the magnitude of the FDR *p*-value. Expression levels of representative top regulated genes within each module and pathway along the TB spectrum is shown in **Supplementary Figure 3**.

M3 showed highly significant enrichment for innate inflammatory pathways well known to be induced in TB ^33, 37, 39^, including IFN, type I IFN, and IFNγ signalling and cytokine signalling (**Figure 3B**). Expression of several canonical IFN-stimulated genes, which are common components of transcriptomic signatures of TB, including FCGR1A, GBP1, 4 and 5, IFIT3, and IFITM3 reflected this progressive increase along the TB spectrum trajectory (**Supplementary Figure 3A**).

M5 showed highly significant enrichment for B cell and B cell receptor signalling pathways (**Figure 3B**) and several canonical B cell genes, such as CD19, CD79A and B, and IGHD, reflecting a pattern of progressive decrease along the TB spectrum trajectory (**Supplementary Figure 3B**).

M4 showed enrichment of pathways associated with translation and peptide chain elongation and included genes encoding ribosomal proteins, such as RPL31, RPL34, RPL7, and RPS7 (**Supplementary Figure 3C**).

No significant enrichment for a highly informative pathway could be identified for M2 or M6 (**Figure 3B**), although progressive decreases along the TB spectrum were observed for M6 genes modulated by signal transduction through the fibroblast growth factor receptor (FGFR), such as CAMK4, CD28, THEM4, and RASGRF2 (**Supplementary Figure 3D**).

### Performance of published transcriptomic TB signatures in asymptomatic TB

Next, we sought to determine how previously published, validated transcriptomic signatures of TB perform as diagnostic biomarkers of asymptomatic TB. We selected a set of signatures that were developed as diagnostic biomarkers for TB (Kaforou24^36^, Maertzdorf4^40^, Sweeney3^41, 42^, XpertHR^43^), prognostic biomarkers for TB (Gliddon3^44^, Penn-Nicholson6^45^, Roe1^46^, and Suliman4^47^), a paediatric TB signature (Thakur9^48^), and a signature of *M. tuberculosis* containment in mice with promising performance in humans (Duffy5^49^). All 10 signatures significantly discriminated between asymptomatic TB and healthy controls in both the HHC Screening and the Community Screening cohorts, with largely similar AUROC values (**Supplementary Figure 4)**. AUROC values were generally higher (AUROCs mostly above 0.75) in the latter (AUROCs mostly below 0.75) than the former cohort. However, it was notable that discriminatory performance for all signatures was better for symptomatic TB than for asymptomatic TB, with the exception of the Duffy5 signature. Analysis of published, validated transcriptomic signatures of TB therefore showed the same pattern as reported above, namely that asymptomatic TB was characterised by less systemic inflammatory signal than symptomatic TB.

### Serum proteomic profiles of asymptomatic TB

We then explored changes in acute response, inflammatory, and immune response serum protein mediators associated with asymptomatic TB, relative to symptomatic TB. Among HHC Screening Cohort participants, serum protein data were available for 28 asymptomatic TB cases, 7 symptomatic TB cases, and 136 controls. Among symptomatic Clinic Triage Cohort participants, serum protein data were available for 206 TB cases and 460 ORD controls, and among Community Screening Cohort participants, protein data were available for 27 asymptomatic TB cases, 6 symptomatic TB cases, and 180 controls (**Supplementary Table 1**).

Overall, diagnostic performance for distinguishing TB from TB-negative controls across community screening cohorts, measured by AUROC, was consistently higher for individuals with symptomatic TB compared with those with asymptomatic TB. This trend was observed in both HHC and Community Screening Cohorts, indicating stronger discriminatory signal in symptomatic disease. Among the top-performing biomarkers were CXCL10, IFN-γ, and IL-6. In the Community HHC Screening Cohort, these markers attained AUROC 0.683 (95%CI 0.583-0.783), 0.545 (95%CI 0.427-0.663), and 0.565 (95%CI 0.460-0.670) in asymptomatic TB, and 0.74 (95%CI 0.546-0.926), 0.720 (95%CI 0.444-0.960), and 0.707 (95%CI 0.472-0.942) in symptomatic TB respectively. In the Community Screening Cohort, the discriminatory power was 0.770 (95%CI 0.695-0.846), 0.674 (95%CI 0.590-0.758), and 0.722 (95%CI 0.638-0.806) in asymptomatic TB, and 0.981 (95%CI 0.965-0.998), 0.820 (95%CI 0.641-1.00), and 0.921 (95%CI 0.879-0.962) in symptomatic TB respectively (**Figure 4A** and **Supplementary Figure 5**). CXCL10 consistently ranked the best in both asymptomatic and symptomatic across both cohorts. Notably, performance metrics were generally higher in the Community than the HHC Screening Cohort, suggesting possible cohort-specific difference in disease severity and/or host response (**Figure 4A** & **B** and **Supplementary Figure 5A** & **B**).

**Figure 4.**
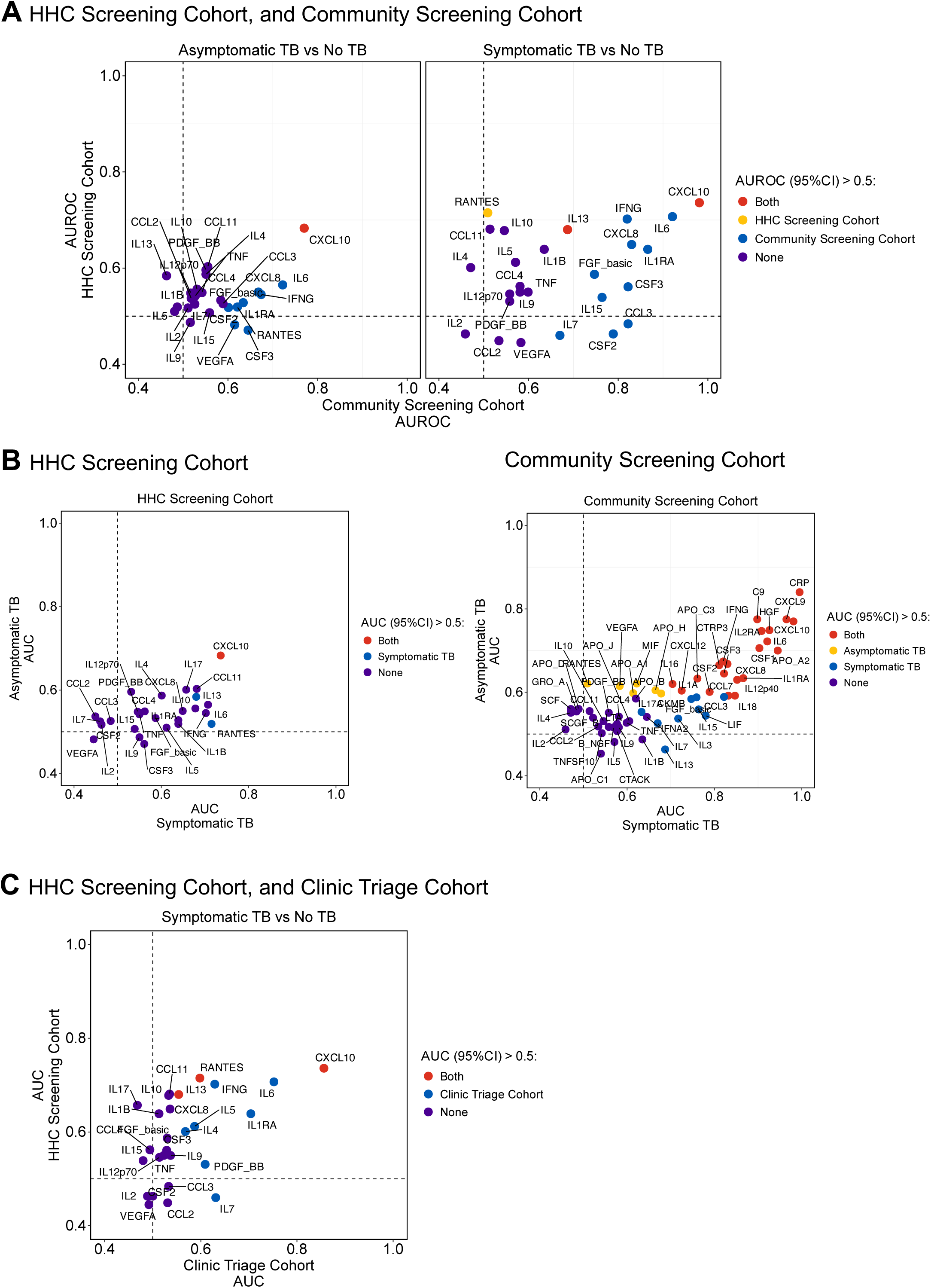
Proteomic biomarkers of asymptomatic TB. Scatter plots summarizing the diagnostic performance (AUROC) of proteomic biomarkers for distinguishing between asymptomatic TB and TB-negative controls in HHC Screening (RePORT SA, Y-axis) and Community Screening (CORTIS, X-axis) participants or between symptomatic TB and from TB-negative controls in Clinic Triage participants. Each point represents a protein, with its position reflecting AUROC values for the relevant comparison. Points are color-coded according to their lower 95% confidence interval (CI) value for the AUROC; biomarkers with lower 95%CI values below 0.5 were considered to display no significant differentiation between the groups (purple), those with lower 95%CI values above 0.5 in only one cohort are in yellow or blue, and those with lower 95%CI values above 0.5 in both cohorts are in red. (**A**) AUROC in asymptomatic TB vs control (left) and symptomatic TB vs control (right) in the Community HHC Screening (Y-axis), and Community Screening (X-axis) cohorts for proteins measured in both cohorts. (**B**) AUROC in symptomatic TB vs controls in Community HHC Screening (Y-axis), and in Clinic Triage participants (X-axis) for protein markers measured in both cohorts. (**C**) AUROC in asymptomatic TB vs control (Y-axis) and symptomatic TB vs control (X-axis) in the Community HHC Screening (left), and in Community Screening (right) cohorts using all proteins measured in the respective cohorts. This visualization highlights both cohort-specific and phenotype-specific performance as well as biomarkers with reproducible diagnostic potential. Data for each protein are in **Supplementary Figure 5**.

We next evaluated these proteins in the Clinic Triage Cohort. A number of proteins showed good diagnostic performance, with the top performing ones including CXCL10 (AUROC 0.856; 95%CI 0.822-0.889%), IL-6 (AUROC 0.752; 95%CI 0.713-0.791), IL-1RA (AUROC 0.704; 95%CI 0.660-0.749), IL-7 (AUROC 0.631; 95%CI 0.586-0.675), and IFNγ (AUROC 0.629; 95%CI 0.583-0.674) (**Figure 4C** and **Supplementary Figure 5C**). Overall, the performance in this cohort was higher than in the asymptomatic and symptomatic cases from the HHC Screening Cohort.

### A transcriptomic signature of asymptomatic TB

Given that previously discovered transcriptomic biomarkers performed poorer for asymptomatic than symptomatic TB, we reasoned that a distinct host response in asymptomatic TB may require a signature that is specific for this manifestation within the TB spectrum. We therefore aimed to discover a signature of asymptomatic TB. To overcome differences in design between the Community HHC Screening (RePORT SA) and Community Screening (CORTIS) cohorts, we combined these cohorts and then partitioned them into discovery (training, 75%) and validation (25%) sets (**Figure 5**). Feature selection and model fitting in the discovery subset resulted in a novel 3-gene signature of asymptomatic TB, comprising *PSTPIP2*, *CARD16,* and *LIMK1*, which discriminated between asymptomatic TB and healthy controls with an AUROC of 83.3% (95% CI 66.5-100%) (**Figure 5A**).

**Figure 5.**
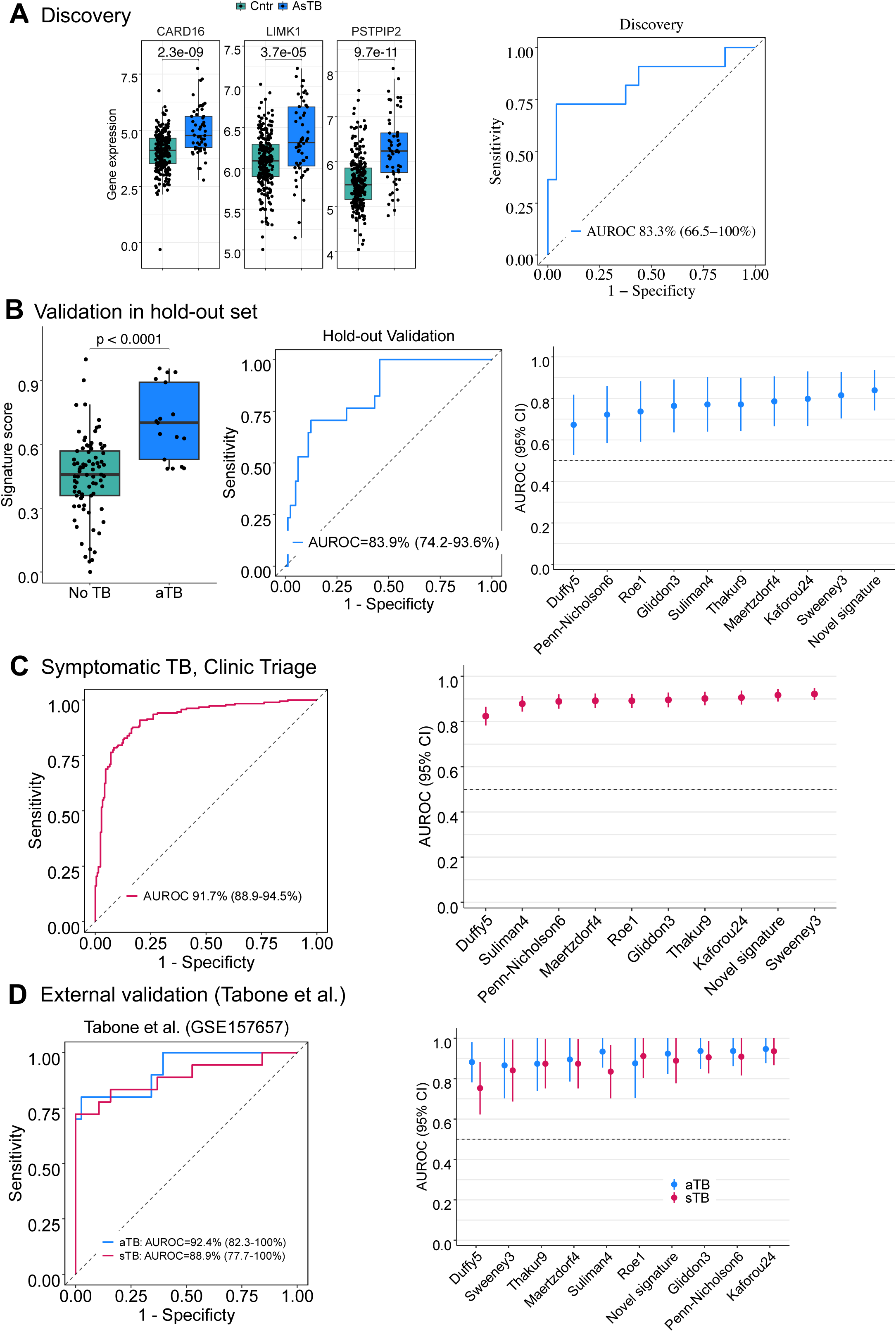
Discovery and validation of a transcriptomic signature of asymptomatic TB. Data from Asymptomatic TB and Control participants from the HHC Screening (RePORT SA) and Community Screening (CORTIS) cohorts were combined and then partitioned into discovery (training, 75%, 21 + 35 asymptomatic TB cases and 135 + 108 healthy controls, respectively) and hold-out validation (25%, 6 + 11 asymptomatic TB cases and 45 + 36 healthy controls, respectively) sets. Feature selection of genes differentiating between asymptomatic TB cases and healthy controls and model fitting into a 3-gene signature of asymptomatic TB, comprising *CARD16, LIMK1* and *PSTPIP2* was performed in the discovery subset. (**A**) Expression levels of genes in the novel asymptomatic TB signature in the discovery set and the receiver-operating characteristic (ROC) curve plot for its performance (model fit) in the discovery set. (**B**) Asymptomatic TB signature scores in asymptomatic TB cases and TB-negative controls in the hold-out validation set. The horizontal lines indicate the median, the boxes the interquartile range (IQR) and the whiskers the range of data with 1.5 x *IQR* from the lower and upper quartiles. The shown *p*-value was computed with the Mann-Whitney U test. The ROC curve shows the performance of the novel signature in the hold-out validation set for differentiating between asymptomatic TB cases and healthy controls. The right-hand plot shows AUROC values (dots) and 95% CIs (error bars) of the novel asymptomatic TB signature as well as a set of previously published TB signatures in the hold-out validation set. (**C**) ROC curve showing the performance of the novel asymptomatic TB signature for differentiating between symptomatic TB and symptomatic TB-negative controls in the Clinic Triage cohort (RePORT SA). The right-hand plot shows AUROC values (dots) and 95% CIs (error bars) of the novel signature and a set of previously published TB signatures in Clinic Triage cohort. (**D**) ROC curves showing the performance of the novel asymptomatic TB signature for differentiating between asymptomatic TB cases (blue) or symptomatic TB cases (red) and controls in the datasets (GSE157657) published by Tabone *et al.*, 2017. The right-hand plot shows AUROC values (dots) and 95% CIs (error bars) of the novel asymptomatic TB signature as well as a set of previously published TB signatures in the GSE157657 dataset.

When applied to the held-out validation set, scores for the novel signature were significantly higher in asymptomatic TB than TB-negative controls; and discriminated between these groups with an AUROC of 83.9% (95% CI 74.2-93.6%, **Figure 5B**). The novel signature nominally had the highest AUROC, but was not significantly better than any of the other 10 transcriptomic signatures. When we analysed diagnostic performance for TB among symptomatic Clinic Triage Cohort participants, AUROC for the novel signature was 91.9% (95% CI 89.2-94.7%), which was also not different to 9 of the 10 published signatures (**Figure 5C**). Finally, we also applied these signatures to a publicly available dataset from HHC in Leicester, UK, described by Tabone *et al*., ^50^, which comprised 38 healthy controls, 7 individuals (10 samples) with “subclinical TB”, defined as having no symptoms but with radiological changes or microbiological evidence of active TB, and 14 individuals with symptomatic, “clinical TB” (microbiologically confirmed). The novel signature, along with 9 of the 10 other signatures discriminated between controls and “subclinical TB” or “clinical TB” with similar performance, as reported for multiple signatures in Tabone et al., ^50^ (**Figure 5D**). Taken together, these results suggest that performance of our novel asymptomatic TB signature was not significantly different to published signatures that were trained for symptomatic TB, progression to incident TB, or TB treatment monitoring.

### Identifying asymptomatic TB sub-types

A critical question is whether there is clustering of features among asymptomatic TB individuals that supports the existence of sub-types that may be associated with clinically-relevant factors, such as infectiousness. Similarity network fusion (SNF) analysis of the full transcriptomic and proteomic datasets from asymptomatic TB cases, combined from the two community screening cohorts, revealed two clusters (**Figure 6A**) by both the rotation cost and Eigen-gap methods. The fully fused network exhibited significant internal stability (ADM=0.03, and APN=0.07; **Supplementary Figure 6**) and median ARI was 90.90% (IQR 82.23-90.91%) across the 500 resampling iterations, each involving a random selection of 80% of the participant samples, in line with established methodological practices^51^.

**Figure 6.**
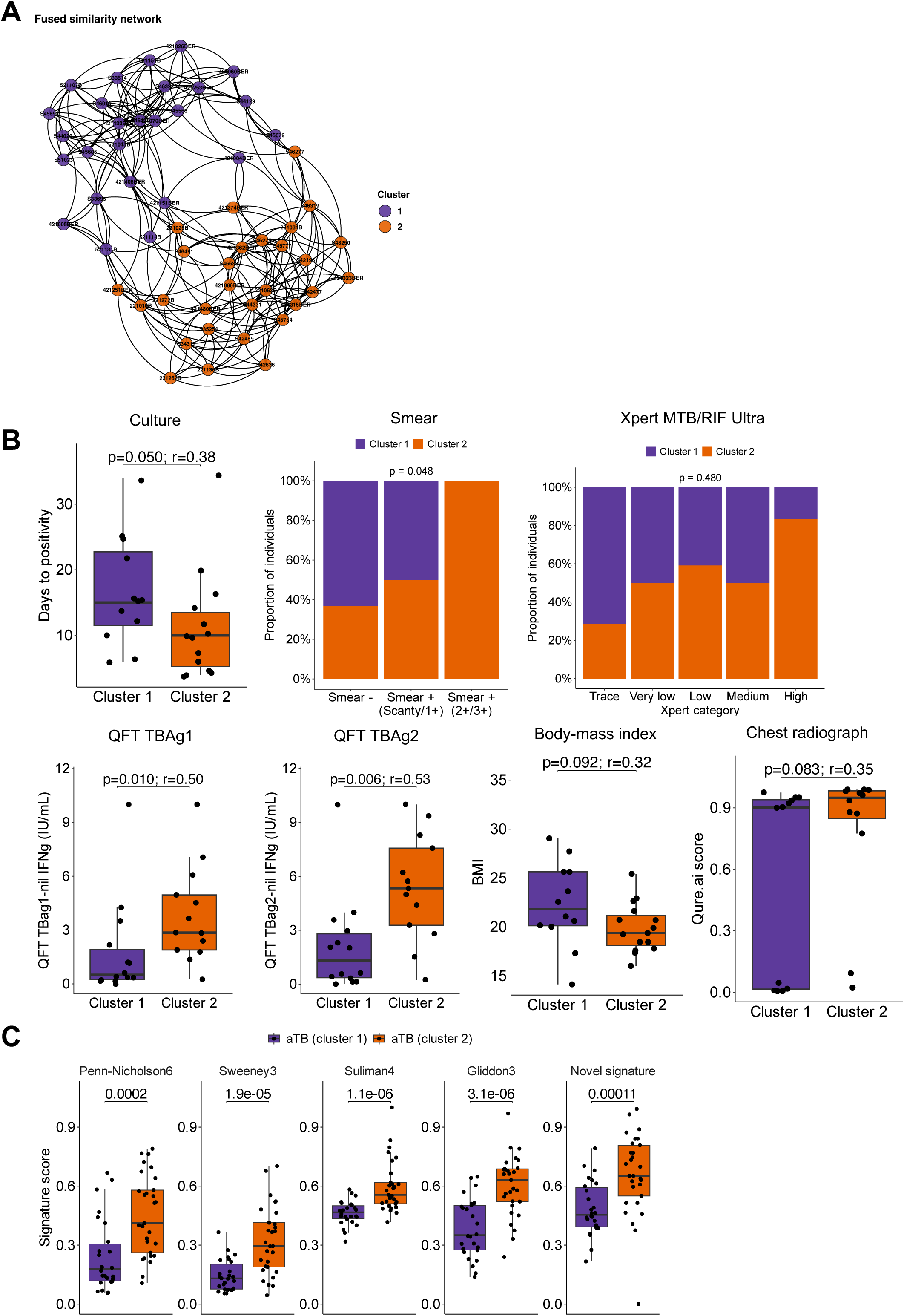
Two asymptomatic TB clusters based on transcriptomic and proteomic data. (**A**) Similarity network fusion (SNF) analysis on the full transcriptomic and proteomic datasets from the HHC and Community Screening Cohorts combined identified two clusters among asymptomatic TB individuals. Stability of these clusters is addressed in **Supplementary Figure 6** (**B**) Sputum bacteriological outcomes, including days to liquid MGIT culture positivity (only those with culture-positive results included), sputum smear microscopy grade and Xpert semiquantitative category (only those with Xpert-positive or trace results included) among asymptomatic TB clusters 1 and 2 individuals in the HHC and Community Screening Cohorts. IFNγ levels from QuantiFERON Gold- Plus assays for TBAg1 or TBAg2, with levels from the Nil condition subtracted, and BMI values for cluster 1 and 2 individuals in the HHC Screening Cohort (RePORT SA). BMI data were only available for the Community Screening Cohort (CORTIS); analysis of smear grade, and IFNγ levels from QuantiFERON Gold-Plus assays, and CXR scores was based on the Community HHC Screening Cohort (RePORT SA) only, whereas culture and Xpert grading was available for both cohorts. (**C**) Transcriptomic signature scores for the indicated TB signatures in cluster 1 and 2 individuals. The shown *p*-values were computed with the Mann-Whitney U test. For box-and-whisker plots horizontal lines indicate the median, the boxes the interquartile range (IQR) and the whiskers the range of data with 1.5 x *IQR* from the lower and upper quartiles.

To explore whether disease severity differed between the two asymptomatic TB clusters, we compared sputum microbiological outcomes. Individuals in cluster 2 had lower time to sputum MGIT culture positivity, a higher proportion of 2+ or 3+ sputum smear grade, and a higher proportion of Xpert semiquantitative high results, all consistent with higher bacterial burden, than individuals in cluster 1 (**Figure 6B**). Cluster 2 individuals also had significantly higher levels of QFT Plus IFN-γ levels, lower BMI, and higher Qure.ai CXR scores, than individuals in cluster 1, further supporting that those in cluster 2 had more severe disease than those in cluster 1.

Finally, we compared selected transcriptomic TB signatures, Penn-Nicholson6, Sweeney3, Suliman4, Gliddon3 and the newly discovered asymptomatic TB signature in the two asymptomatic TB clusters. Signature scores for all these signatures were significantly higher in cluster 2 individuals than those in cluster 1 (**Figure 6C**), suggesting higher systemic inflammatory responses in cluster 2. Interestingly, cluster 1 had similar Sweeney3 and Duffy5^49^ signature scores to No-TB controls, while cluster 2 had similar signature scores to symptomatic TB cases (**Supplementary Figure 7**) The same was observed for granulocyte/lymphocyte ratio, a marker of systemic inflammation. Finally, a similar pattern was seen in analyses of gene expression and protein markers that informed the clustering, all showing that cluster 1 individuals had lower levels of inflammatory markers, while cluster 2 comprised individuals with more inflammation, at levels similar to those observed in Community Screening individuals with symptomatic TB (**Supplementary Figure 8**).

## Discussion

We characterised blood transcriptomic and proteomic profiles in asymptomatic TB detected by household and community screening, compared to symptomatic TB detected by facility-based triage, in contemporaneous cohorts using standardized approaches. Our results show that systemic inflammatory profiles among persons with asymptomatic TB, including proteomic and transcriptomic signatures and MDH were highly heterogenous; and of significantly lower magnitude than inflammatory profiles among individuals with symptomatic TB, whether detected in the community or presenting to clinics. These findings are consistent with previous findings from RePORT SA, showing that CRP levels, yield of CXR screening, and sputum bacillary load were significantly lower in asymptomatic TB than is typically observed in symptomatic TB ^6^.

Most TB diagnostic studies enrol symptomatic patients who seek care at health facilities, typically with more advanced disease than is identified in community screening studies. Therefore, the poor performance of host response biomarkers for asymptomatic TB, relative to symptomatic TB, has important implications for feasibility of biomarker-based screening approaches to identify individuals with undiagnosed TB in community settings. These data address two fundamental questions that may be critical for TB control: first, whether asymptomatic differs from symptomatic TB in biology and severity (it does); and second, whether biomarkers developed and validated for symptomatic TB in triage settings perform adequately for detection of asymptomatic TB in community settings (they do not). The latter question is highly relevant, since the World Health Organization (WHO) recently combined triage and screening test guidelines under a single Target Product Profile^18^. Yet, our findings suggest that biomarkers developed for facility-based triage of symptomatic TB cannot simply be re-purposed for community-based screening of asymptomatic TB without further optimization or acceptance of lower performance^6^.

Application of published transcriptomic signatures of TB disease performed sub-optimally to distinguish between asymptomatic TB and healthy controls. We therefore set out to discover a novel, parsimonious transcriptomic signature of asymptomatic TB. This signature did not significantly outperform published transcriptomic signatures of TB in a hold-out validation cohort, nor in a previously published cohort of asymptomatic TB^50^.

GSEA and WGCNA of RNA-seq data suggested that similar biological pathways and processes were perturbed in asymptomatic TB, albeit to a lesser degree than the well-described perturbations in symptomatic TB ^33, 37, 39, 45, 50, 52, 53^. The most prominent pathways found to be induced during asymptomatic TB were IFN responses, complement activation, inflammation, and myeloid cell activation, with peripheral depletion of lymphoid cells such as B cells, T cells, and cytotoxic lymphocytes, including NK cells. These pathways are consistent with those reported for symptomatic TB, as well as studies of TB progression, and thus our findings do not suggest that asymptomatic TB is a distinct phenotype of TB. Rather, these findings, along with MDH, WGCNA, pathway analyses and proxy measures of sputum bacillary load, indicate that asymptomatic TB occupies an intermediate state within the TB spectrum, between early disease (seen in those who ultimately progress to incident TB) and those with more severe, symptomatic disease ^39, 45, 47, 52, 54^.

A major finding of our study is that clustering analysis of transcriptomic and proteomic data from individuals with asymptomatic TB segregated into two stable clusters that may represent sub-types (**Supplementary Method**). These two clusters differed in sputum bacillary burden, CXR extent of disease, IGRA responses, and transcriptomic signature scores, consistent with differences in extent or severity of asymptomatic disease. Important questions include whether those with more severe pulmonary disease are more likely to progress to symptomatic TB, or if they may be more infectious. Recent studies showed that symptoms, such as coughing, are not required for aerosol release of *M. tuberculosis* ^8, 9^, implying that individuals with asymptomatic TB may contribute to transmission of the pathogen.

Our finding of marked heterogeneity among those with asymptomatic TB is consistent with a PET-CT imaging study in close contacts of TB cases with rifampicin-resistant TB, which also showed a large range of pulmonary and other thoracic manifestations ^19^. A major finding was a clear distinction between individuals with PET/CT-active and PET/CT-inactive pulmonary lesions, where presence of PET/CT-active lesions was associated with more than 20-fold increased risk of progressing to TB disease and self presentation with symptoms to the health services. It is possible that the two clusters we identified represent two sub-types with relatively metabolically active vs quiescent pulmonary or lymphatic lesions. A number of longitudinal studies that compared TB progressors with non-progressors have shown higher transcriptomic and proteomic signature scores, as well as activation of *M. tuberculosis*-specific T cells among progressors ^4, 5, 6, 17, 39, 44, 45, 47, 52, 54, 55^. We speculate that these sub-types of asymptomatic TB might reflect pre-symptomatic progressors and others with stable, chronic, or even regressing pathology.

Future studies with longitudinal follow-up of those with asymptomatic TB is necessary to discern between those who may progress to symptomatic TB and those who may remain asymptomatic, or those who may regress to clear mycobacteria from their respiratory tract. Such studies should include detection of *M. tuberculosis*-containing bioaerosol release to inform infectiousness of asymptomatic disease ^8, 9^. Such a study raises an ethical dilemma about whether people with bacterially-confirmed TB should receive multidrug treatment, even if they appear well and reluctant to partake in a 4-6 month regimen.

Our study has several limitations. Although the HHC and Community Screening Cohorts were large, the yield of asymptomatic TB cases was relatively low. We cannot rule out that we have under-ascertained characteristics of asymptomatic TB; and that more sub-types might exist. We found that transcriptomic and proteomic differences between asymptomatic TB cases and controls were higher magnitude in the CORTIS, compared to the RePORT SA cohorts, suggesting either laboratory or assay differences, or differences in study design, study population, duration of disease, and/or time since exposure, which may be relevant to HHC cohorts. However, procedures and assays were performed in the same SATVI laboratory; and sample processing was highly consistent between the CORTIS and RePORT SA studies, ruling out technical differences as a major factor. The most likely explanation is that the CORTIS study, which pre-screened the study population to enrich for RISK11 signature-positive individuals, selected for those with more systemic inflammation. However, this factor would tend to minimize, rather than exaggerate the differences between asymptomatic and symptomatic TB observed in this study.

Regardless, to our knowledge this is the largest study of host blood proteomic and transcriptomic biomarkers in asymptomatic TB; and standardized data collection in contemporaneous cohorts allows direct comparison with symptomatic TB. Our findings suggest that biomarker approaches based on blood inflammatory markers are unlikely to be effective for community screening for asymptomatic TB without further optimization. Further, the clear differences in inflammation and severity of disease between individuals with symptomatic TB presenting to health facilities and those with asymptomatic TB detected in the community suggest that fundamental differences in biology should be taken into account when developing policy for TB screening and triage tests^18^.

## Author contributions

TJS, TRS, and MH conceived the idea, raised funds, and provided the resources. GW, TJS, TRS, and MH wrote the RePORT-South Africa study protocol and provided study oversight. TJS, AF-G, GW, KN, GC, and MH designed the CORTIS-01 study and provided study oversight. MT, TM, STM, FN, CI, WB, AH, KN, KD, SJ, TP, NM, AL, BF and GC were responsible for site-level activities, including recruitment, clinical management, and data collection. S.N processed and analysed chest Xrays data. A.F.G provided analytical support for the CORTIS-01 study. D.A designed machine learning pipeline and conducted data analyses. DTA, VR, and NNC processed and analysed Bio-Plex data. KHa provided operational support and project management. HM, AK, FM, RP, KHl, KS, and YFvdH cleaned and verified the underlying data. TJS, HM, TS, and MH analysed data and interpreted results. DA, SCM, MH and TJS drafted the manuscript.

Members of the RePORT SA team, and CORTIS study team performed clinical studies and processed samples. All authors reviewed, edited, and approved the final version of the manuscript submitted.

## Data availability

Deidentified, processed gene expression data and raw fast data will be deposited on GEO repository. Luminex data is available upon request from the corresponding author.

