## Supplemental material for "Inflammatory Biomarkers of Asymptomatic and Symptomatic Tuberculosis"

### SUPPLEMENTARY INFORMATION

#### *RePORT South Africa Study Team

| Name | Affiliation |
|---|---|
| Pattamukkil Abraham | Perinatal HIV Research Unit, University of Witwatersrand |
| Thakiera Allie | South African Tuberculosis Vaccine Initiative, University of Cape Town |
| Cynthia Baard | Department of Paediatrics and Child Health, University of Cape Town |
| Zainab Baig | Africa Health Research Institute |
| John Belisle | Colorado State University |
| Nicole Bilek | South African Tuberculosis Vaccine Initiative, University of Cape Town |
| Gerard Cangelosi | University of Washington |
| ACE Carstens | Department of Biomedical Sciences, Stellenbosch University |
| Kevyna Chetty | Africa Health Research Institute |
| Yolundi Cloete | South African Tuberculosis Vaccine Initiative, University of Cape Town |
| Marwou de Kock | South African Tuberculosis Vaccine Initiative, University of Cape Town |
| Gareta Dickman | Africa Health Research Institute |
| Charity Dire | Perinatal HIV Research Unit, University of Witwatersrand |
| Karen Dobos | Colorado State University |
| Stephany Norah Duda | Vanderbilt Tuberculosis Center, Vanderbilt University |
| Mzwandile Erasmus | South African Tuberculosis Vaccine Initiative, University of Cape Town |
| Marina Cruvinel Figueiredo | Vanderbilt Tuberculosis Center, Vanderbilt University |
| Marika Flinn | Department of Biomedical Sciences, Stellenbosch University |
| Travis Harris | Vanderbilt Tuberculosis Center, Vanderbilt University |
| Ryan Johnson | University of Cape Town |

| | |
|---|---|
| Farina Karim | Africa Health Research Institute |
| Masooda Kaskar | South African Tuberculosis Vaccine Initiative, University of Cape Town |
| Nobulumko Khomba | South African Tuberculosis Vaccine Initiative, University of Cape Town |
| Thandeka Khoza | Africa Health Research Institute |
| Léanie Kleynhans | Department of Biomedical Sciences, Stellenbosch University; Mater Research Institute – The University of Queensland |
| Tahira Kootbodien | University of Cape Town |
| Andrea Kotze | University of Cape Town Lung Institute |
| Belinda A Kriel | Department of Biomedical Sciences, Stellenbosch University |
| Ané Kruger | Department of Biomedical Sciences, Stellenbosch University |
| Lorraine Lichakane | Perinatal HIV Research Unit, University of Witwatersrand |
| Ilze Louw | Department of Biomedical Sciences, Stellenbosch University |
| Angelique Luabeya | South African Tuberculosis Vaccine Initiative, University of Cape Town |
| Candice MacDonald | Department of Biomedical Sciences, Stellenbosch University |
| Lindiwe Madziwa | Africa Health Research Institute |
| Lebohang Makhethe | South African Tuberculosis Vaccine Initiative, University of Cape Town |
| Sandisiwe Mangali | South African Tuberculosis Vaccine Initiative, University of Cape Town |
| Linda Mbuthini | University of Cape Town |
| Carolina Mehaffy | Colorado State University |
| Thabang Moloja | Perinatal HIV Research Unit, University of Witwatersrand |
| Angelique Mouton | South African Tuberculosis Vaccine Initiative, University of Cape Town |
| Evans Muchiri | South African Tuberculosis Vaccine Initiative, University of Cape Town |
| Mbusiseni Ngema | Perinatal HIV Research Unit, University of Witwatersrand |
| Hlengiwe Nkambule | South African Tuberculosis Vaccine Initiative, University of Cape Town |
| Onke Nombida | South African Tuberculosis Vaccine Initiative, University of Cape Town |
| Alaina Olson | University of Washington |
| Fajwa Opperman | South African Tuberculosis Vaccine Initiative, University of Cape Town |
| Gregory Ording-Jespersen | Africa Health Research Institute |
| Kennedy Otwombe | Perinatal HIV Research Unit, University of Witwatersrand |
| Will Ramsay | Vanderbilt Tuberculosis Center, Vanderbilt University |
| Tracy Richardson | Department of Biomedical Sciences, Stellenbosch University |
| Carmen Segelaar | South African Tuberculosis Vaccine Initiative, University of Cape Town |
| Jane Shaw | Department of Biomedical Sciences, Stellenbosch University |
| Kimberly Shelton | Colorado State University |
| Justin Shenje | South African Tuberculosis Vaccine Initiative, University of Cape Town |
| Theresa Smit | Africa Health Research Institute |

| | |
|---|---|
| Bronwyn Smith | Department of Biomedical Sciences, Stellenbosch University |
| Candice Snyders | Department of Biomedical Sciences, Stellenbosch University |
| Marcia Steyn | South African Tuberculosis Vaccine Initiative, University of Cape Town |
| Sara Suliman | University of California San Francisco |
| Floris Swanepoel | Perinatal HIV Research Unit, University of Witwatersrand |
| Lorraine Thobakgale | Perinatal HIV Research Unit, University of Witwatersrand |
| Susanne Tonsing | Department of Biomedical Sciences, Stellenbosch University |
| Nicolette Tredoux | South African Tuberculosis Vaccine Initiative, University of Cape Town |
| Megan Turner | Vanderbilt Tuberculosis Center, Vanderbilt University |
| Petrus Tyambetyu | South African Tuberculosis Vaccine Initiative, University of Cape Town |
| Habibullah Valley | South African Tuberculosis Vaccine Initiative, University of Cape Town |
| Lyle van de Berg | University of Pretoria |
| Gian van der Spuy | Department of Biomedical Sciences, Stellenbosch University |
| Ilana R van Rensburg | Department of Biomedical Sciences, Stellenbosch University |
| Johanna E van Rooyen | South African Tuberculosis Vaccine Initiative, University of Cape Town |
| Hilary Vansell Riley | Vanderbilt Tuberculosis Center, Vanderbilt University |
| Ashley Veldsman | South African Tuberculosis Vaccine Initiative, University of Cape Town |
| Lindsay Wilson | University of Cape Town |
| Rachel C Wood | University of Washington |
| Lesley Workman | Department of Paediatrics and Child Health, University of Cape Town |
| Heather Zar | Department of Paediatrics and Child Health, University of Cape Town |

***CORTIS Study Team**

| First name | Surname | Affiliation |
|---|---|---|
| Laudicia Tshenolo | Bontsi | The Aurum Institute |
| Obakeng Peter | Booi | The Aurum Institute |
| Mari Cathrin | Botha | The Aurum Institute |
| Kgomotso Violet | Chauke | The Aurum Institute |
| Mooketsi Theophillius | Cwaile | The Aurum Institute |
| Isabella Johanna | Davies | The Aurum Institute |
| Emilia | De Klerk | The Aurum Institute |
| Blanchard Mbay | Iyemosolo | The Aurum Institute |
| James Michael | Jeleni | The Aurum Institute |
| Christian Mabika | Kasongo | The Aurum Institute |
| Christian Mabika | Kasongo | The Aurum Institute |
| Sebaetseng Jeanette | Kekana | The Aurum Institute |
| Lucky Sipho | Khoza | The Aurum Institute |

| | | |
|---|---|---|
| Gloria Keitumetse | Kolobe | The Aurum Institute |
| Lerato Julia | Lekagane | The Aurum Institute |
| Sheiley Christina | Lekotloane | The Aurum Institute |
| Ilze Jeanette | Louw | The Aurum Institute |
| Sarah Teboso | Lusale | The Aurum Institute |
| Perfect Tiisetso | Maatjie | The Aurum Institute |
| Kamogelo Fortunate | Mabena | The Aurum Institute |
| Johanna Thapelo | Madikwe | The Aurum Institute |
| Octavia Mahkosazana | Madikwe | The Aurum Institute |
| Rapontwana Letlhogonolo | Maebana | The Aurum Institute |
| Malobisa Sylvester | Magwasha | The Aurum Institute |
| Vutlhari-I-Vunhenha Fairlord | Manzini | The Aurum Institute |
| Isholedi Samuel | Maroele | The Aurum Institute |
| Omphile Petunia | Masibi | The Aurum Institute |
| July Rocky | Mathabanzini | The Aurum Institute |
| Tendamudzimu Ivan | Mathode | The Aurum Institute |
| Ellen Ditaba | Matsane | The Aurum Institute |
| Lungile | Mbata | The Aurum Institute |
| Nyasha Karen | Mhandire | The Aurum Institute |
| Thembisiwe | Miga | The Aurum Institute |
| Nosisa Charity Thandeka | Mkhize | The Aurum Institute |
| Caroline | Mkhokho | The Aurum Institute |
| Neo Hilda | Mkwalase | The Aurum Institute |
| Nondzakazi | Mnqonywa | The Aurum Institute |
| Brenda Matshidiso | Modisaotsile | The Aurum Institute |
| Patricia Pakiso | Mokgetsengoane | The Aurum Institute |
| Selemeng Matseliso Carol | Mokone | The Aurum Institute |
| Kegomoditswe Magdeline | Molatlhegi | The Aurum Institute |
| Thuso Andrew | Molefe | The Aurum Institute |
| Motlatsi Evelyn | Molotsi | The Aurum Institute |
| Tebogo Edwin | Montwedi | The Aurum Institute |
| Boikanyo Dinah | Monyemangene | The Aurum Institute |
| Hellen Mokopi | Mooketsi | The Aurum Institute |
| Tshplpfelo Mapula | Mosito | The Aurum Institute |
| Ireen Lesebang | Mosweu | The Aurum Institute |
| Banyana Olga | Motlagomang | The Aurum Institute |
| Funeka Nomvula | Mthembu | The Aurum Institute |
| Themba | Phakathi | The Aurum Institute |
| Mapule Ozma | Phatshwane | The Aurum Institute |
| Victor Kgothatso | Rameetse | The Aurum Institute |

| | | |
|---|---|---|
| Kelebogile Magdeline | Segaetsho | The Aurum Institute |
| Ni Ni | Sein | The Aurum Institute |
| Melissa Neo | Senne | The Aurum Institute |
| Sifiso Cornelius | Shezi | The Aurum Institute |
| Zona | Sithetho | The Aurum Institute |
| Bongiwe | Stofile | The Aurum Institute |
| Mando Mmakhora | Thaba | The Aurum Institute |
| Lethabo Collen | Theko | The Aurum Institute |
| Dimakatso Sylvia | Tsagae | The Aurum Institute |
| Dhineshree | Govender | Centre for the AIDS Programme of Research in South Africa (CAPRISA) |
| Tilagavathy | Chinappa | Centre for the AIDS Programme of Research in South Africa (CAPRISA) |
| Mbali Ignatia | Zulu | Centre for the AIDS Programme of Research in South Africa (CAPRISA) |
| Nonhle Bridgette | Maphanga | Centre for the AIDS Programme of Research in South Africa (CAPRISA) |
| Senzo Ralph | Hlathi | Centre for the AIDS Programme of Research in South Africa (CAPRISA) |
| Goodness Khanyisile | Gumede | Centre for the AIDS Programme of Research in South Africa (CAPRISA) |
| Thandiwe Yvonne | Shezi | Centre for the AIDS Programme of Research in South Africa (CAPRISA) |
| Jabulisiwe Lethabo | Maphanga | Centre for the AIDS Programme of Research in South Africa (CAPRISA) |
| Zandile Patrica | Jali | Centre for the AIDS Programme of Research in South Africa (CAPRISA) |
| Thobelani | Cwele | Centre for the AIDS Programme of Research in South Africa (CAPRISA) |
| Nonhlanhla Zanele Elsie | Gwamanda | Centre for the AIDS Programme of Research in South Africa (CAPRISA) |
| Celaphiwe | Dlamini | Centre for the AIDS Programme of Research in South Africa (CAPRISA) |
| Zibuyile Phindile Penlee | Sing | Centre for the AIDS Programme of Research in South Africa (CAPRISA) |
| Ntombozuko Gloria | Ntanjana | Centre for the AIDS Programme of Research in South Africa (CAPRISA) |
| Sphelele Simo | Nzimande | Centre for the AIDS Programme of Research in South Africa (CAPRISA) |
| Siyabonga | Mbatha | Centre for the AIDS Programme of Research in South Africa (CAPRISA) |
| Bhavna | Maharaj | Centre for the AIDS Programme of Research in South Africa (CAPRISA) |
| Atika | Moosa | Centre for the AIDS Programme of Research in South Africa (CAPRISA) |
| Cara-Mia | Corris | Centre for the AIDS Programme of Research in South Africa (CAPRISA) |
| Razia Hassan | Moosa | Centre for the AIDS Programme of Research in South Africa (CAPRISA) |
| Fazlin | Kafaar | South African Tuberculosis Vaccine Initiative, University of Cape Town |
| Marwou | De Kock | South African Tuberculosis Vaccine Initiative, University of Cape Town |

| | | |
|---|---|---|
| Hennie | Geldenhuys | South African Tuberculosis Vaccine Initiative, University of Cape Town |
| Angelique Kany Kany | Luabeya | South African Tuberculosis Vaccine Initiative, University of Cape Town |
| Justin | Shenje | South African Tuberculosis Vaccine Initiative, University of Cape Town |
| Natasja | Botes | South African Tuberculosis Vaccine Initiative, University of Cape Town |
| Susan | Rossouw | South African Tuberculosis Vaccine Initiative, University of Cape Town |
| Hadn | Africa | South African Tuberculosis Vaccine Initiative, University of Cape Town |
| Bongani | Diamond | South African Tuberculosis Vaccine Initiative, University of Cape Town |
| Samentra | Braaf | South African Tuberculosis Vaccine Initiative, University of Cape Town |
| Sonia | Stryers | South African Tuberculosis Vaccine Initiative, University of Cape Town |
| Alida | Carstens | South African Tuberculosis Vaccine Initiative, University of Cape Town |
| Ruwiyda | Jansen | South African Tuberculosis Vaccine Initiative, University of Cape Town |
| Simbarashe | Mabwe | South African Tuberculosis Vaccine Initiative, University of Cape Town |
| Roxane | Herling | South African Tuberculosis Vaccine Initiative, University of Cape Town |
| Ashley | Veldsman | South African Tuberculosis Vaccine Initiative, University of Cape Town |
| Lebohgang | Makhete | South African Tuberculosis Vaccine Initiative, University of Cape Town |
| Marcia | Steyn | South African Tuberculosis Vaccine Initiative, University of Cape Town |
| Sivuyile | Buhlungu | South African Tuberculosis Vaccine Initiative, University of Cape Town |
| Margareth | Erasmus | South African Tuberculosis Vaccine Initiative, University of Cape Town |
| Ilse | Davids | South African Tuberculosis Vaccine Initiative, University of Cape Town |
| Patiswa | Plaatjie | South African Tuberculosis Vaccine Initiative, University of Cape Town |
| Alessandro | Companie | South African Tuberculosis Vaccine Initiative, University of Cape Town |
| Frances | Ratangee | South African Tuberculosis Vaccine Initiative, University of Cape Town |
| Helen | Veldtsman | South African Tuberculosis Vaccine Initiative, University of Cape Town |
| Christel | Petersen | South African Tuberculosis Vaccine Initiative, University of Cape Town |
| Charmaine | Abrahams | South African Tuberculosis Vaccine Initiative, University of Cape Town |
| Miriam | Moses | South African Tuberculosis Vaccine Initiative, University of Cape Town |
| Xoliswa | Kelepu | South African Tuberculosis Vaccine Initiative, University of Cape Town |
| Yolande | Gregg | South African Tuberculosis Vaccine Initiative, University of Cape Town |

| | | |
|---|---|---|
| Liticia | Swanepoel | South African Tuberculosis Vaccine Initiative, University of Cape Town |
| Nomsitho | Magawu | South African Tuberculosis Vaccine Initiative, University of Cape Town |
| Nompumelelo | Cetywayo | South African Tuberculosis Vaccine Initiative, University of Cape Town |
| Lauren | Mactavie | South African Tuberculosis Vaccine Initiative, University of Cape Town |
| Habibullah | Valley | South African Tuberculosis Vaccine Initiative, University of Cape Town |
| Elizabeth | Filander | South African Tuberculosis Vaccine Initiative, University of Cape Town |
| Nambitha | Nqakala | South African Tuberculosis Vaccine Initiative, University of Cape Town |
| Angelique | Mouton | South African Tuberculosis Vaccine Initiative, University of Cape Town |
| Fajwa | Opperman | South African Tuberculosis Vaccine Initiative, University of Cape Town |
| Elma | Van Rooyen | South African Tuberculosis Vaccine Initiative, University of Cape Town |
| Petrus | Tyambetyu | South African Tuberculosis Vaccine Initiative, University of Cape Town |
| Elizna | Maasdorp | DST/NRF Centre of Excellence for Biomedical TB Research and SAMRC Centre for TB Research, Stellenbosch University |
| Justine | Khoury | DST/NRF Centre of Excellence for Biomedical TB Research and SAMRC Centre for TB Research, Stellenbosch University |
| Belinda | Kriel | DST/NRF Centre of Excellence for Biomedical TB Research and SAMRC Centre for TB Research, Stellenbosch University |
| Bronwyn | Smith | DST/NRF Centre of Excellence for Biomedical TB Research and SAMRC Centre for TB Research, Stellenbosch University |
| Liesel | Muller | DST/NRF Centre of Excellence for Biomedical TB Research and SAMRC Centre for TB Research, Stellenbosch University |
| Susanne | Tonsing | DST/NRF Centre of Excellence for Biomedical TB Research and SAMRC Centre for TB Research, Stellenbosch University |
| Andre | Loxton | DST/NRF Centre of Excellence for Biomedical TB Research and SAMRC Centre for TB Research, Stellenbosch University |
| Andriette | Hiemstra | DST/NRF Centre of Excellence for Biomedical TB Research and SAMRC Centre for TB Research, Stellenbosch University |
| Petri | Ahlers | DST/NRF Centre of Excellence for Biomedical TB Research and SAMRC Centre for TB Research, Stellenbosch University |
| Marika | Flinn | DST/NRF Centre of Excellence for Biomedical TB Research and SAMRC Centre for TB Research, Stellenbosch University |
| Eva | Chung | Vaccine and Infectious Disease Division, Fred Hutchinson Cancer Research Center |
| Michelle | Chung | Vaccine and Infectious Disease Division, Fred Hutchinson Cancer Research Center |

| | | |
|---|---|---|
| Alicia | Sato | Vaccine and Infectious Disease Division, Fred Hutchinson Cancer Research Center |

### Methods

*Machine Learning for Biomarker Discovery*

To identify a novel host transcriptomic signature of asymptomatic TB, we applied an in-house machine learning pipeline—OmicScan (https://github.com/SATVILab/OmicScan)—to analyse the combined dataset of 397 samples, comprising 46 asymptomatic TB and 144 controls from the HHC Screening and 27 asymptomatic TB and 180 controls from the Community Screening cohorts. We split the combined dataset into a signature discovery set (75%) and a hold-out validation set (25%), and then further partitioned the discovery set into training (20%) and test (20%) sets. We employed stratified sampling to ensure that the splits obtained were not imbalanced with respect to cohort representation and the proportion of the class labels. Each set was independently batch-corrected using ComBat-Seq[1].

The OmicScan pipeline included gene prefiltering to remove noisy/uninformative genes, gene ranking to retain the top outcome-associated genes, and derivation of the parsimonious biomarker model. For gene prefiltering, we applied (1) the t-test with Benjamini-Hochberg procedure to control the false discovery rate (FDR) and retained genes with adjusted p-value (FDR) $<0.05$, and (2) Hedge's g as the measure of effect size (ES) and retained genes with an ES greater than 0.3. Next, we applied four machine learning (ML) algorithms—Support Vector Machine (SVM), Random Forest (RF), Gradient Boosting (XGBoost) and K-Nearest Neighbour (KNN)—to train predictive models and subsequently evaluated their performance on the discovery test set. We used a "g-score", a weighted combination of SHAP (Shapley Additive exPlanations) value and permutation importance score, to rank genes from each ML model based on their contribution to the predictive model. Permutation importance provides a model-agnostic estimate of feature relevance by measuring the decrease in model performance when a feature's value is randomly shuffled. This captured the global impact of each gene on model accuracy[2]. In contrast, SHAP values offered a local interpretability framework quantifying the contribution of each gene to individual predictions based on cooperative game theory[3]. Together, the g-score determined a weighted usefulness of the gene.

Finally, we performed a greedy forward search on the top 10 ranked genes to derive the most predictive, parsimonious combination of genes for diagnostic application. We assessed the diagnostic power of every possible combination of the genes by calculating signature scores of each combination and the corresponding area under the receiver operating characteristic curve (AUROC). For gene combinations containing both up- and down-regulated genes, the signature score was calculated by subtracting the mean expression of the down-regulated genes from that of the up-regulated genes. In contrast, for combinations in which all genes were regulated in the same direction, the signature score was derived using the geometric mean of the expression levels of the associated genes.

*Integration of transcriptomic and proteomic data to discover clusters*

We sought to integrate the abundance of serum proteins and blood RNA-seq datasets from asymptomatic TB cases from the two community screening cohorts to understand the heterogeneity amongst this TB phenotype. We employed Similarity Network Fusion (SNF) in SNFTool[4] to integrate transcriptomic and proteomic data modalities, allowing unsupervised identification of molecular subgroups within the asymptomatic TB dataset. Optimal cluster numbers were identified (2 to 8 clusters) by eigen-gap and rotation cost methods[4].

Given the high heterogeneity and dimensionality of the multiomic data, evaluating cluster validity was essential to minimize the risk of identifying spurious clusters that may arise from algorithmic bias. After identifying subclusters, we utilized the *clValid* R package[5] to assess cluster stability and validity through several metrics, including the average proportion of non-overlap (APN), the average distance between means (ADM), and the adjusted Rand Index (ARI). The APN quantified the extent to which observations changed cluster membership when a subset of features is systematically removed. Specifically, it calculated the average proportion of observations that were assigned to different clusters in a perturbed dataset compared to the original clustering. APN values ranged from 0 to 1, with lower values indicating greater cluster stability under feature perturbation. ADM evaluated the average Euclidean distance between cluster centroids obtained from the original dataset and those obtained after feature removal. ADM values ranged from 0 to 1, with smaller values suggesting that the overall cluster remained consistent across perturbations. ARI, a statistic ranging from 0 (no agreement) to 1 (perfect concordance) measured similarity between two clustering assignments (original and perturbed). Cluster stability was evaluated by performing 500 resampling iterations, each involving a random selection of 80% of the participant samples, in line with established methodological practices[6]. To further examine the relationship between alternative sub-clustering results, we conducted chi-square tests to assess their statistical independence.

### Supplementary Figures

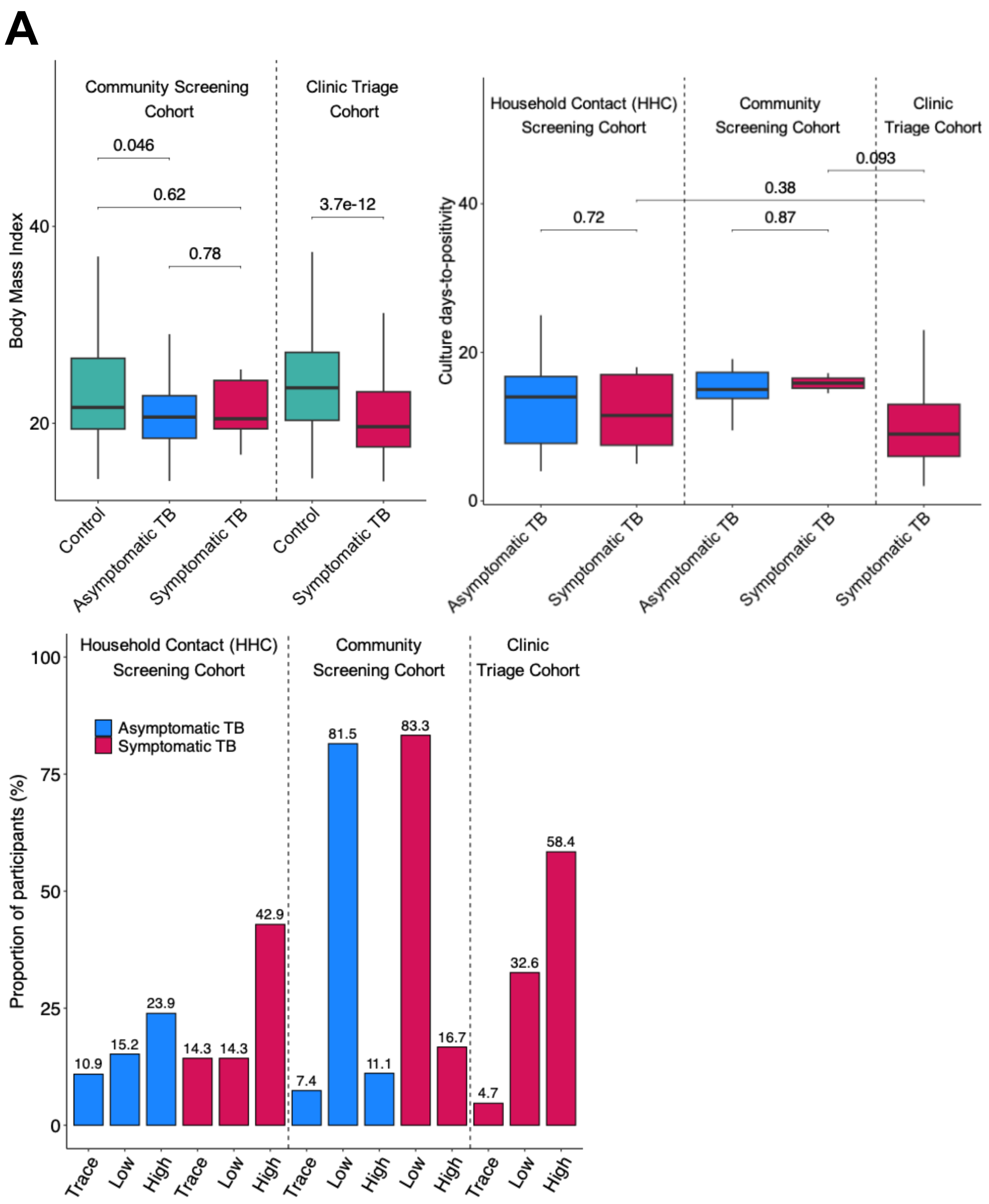


**Supplementary Figure S1. Distribution of clinical and microbiological characteristics of participants in the cohorts.** (**A**) (Top left) Boxplot showing the distribution of body mass index (BMI) by clinical outcome in the Clinic Triage and the HHC Screening cohorts. (Top right) Boxplot showing time to sputum culture positivity (in days) for culture-diagnosed prevalent TB cases in the Clinic Triage, HHC Screening, and the Community Screening cohorts. (Bottom) Bar chart showing the relative proportions of individuals in each Xpert semiquantitative category among asymptomatic TB, and symptomatic TB cases in the cohorts. Proportions were calculated relative to the total number of individuals with each TB outcome, for those with available results. The category "Low" includes 'low' and 'very low', and "High" includes "medium" and "high" Xpert MTB/RIF Ultra categories.

**Supplementary Table 1**. Distributions of age, sex, and HIV status in participants from the HHC Screening Cohort (top), Community Screening Cohort (middle) and Clinic Triage Cohort (bottom).

**HHC Screening Cohort**

| | No TB (N=180) | Asymptomatic TB (N=27) | Symptomatic TB (N=6) | Total (N=231) |
|---|---|---|---|---|
| Age | | | | |
| Median | 28.00 | 32.00 | 26.00 | 28.00 |
| Q1, Q3 | 23.00, 36.25 | 25.00, 36.50 | 24.25, 27.75 | 23.00, 36.00 |
| Sex | | | | |
| Female | 71 (39.4%) | 11 (40.7%) | 4 (66.7%) | 93 (40.3%) |
| Male | 109 (60.6%) | 16 (59.3%) | 2 (33.3%) | 138 (59.7%) |
| BMI | | | | |
| Median | 21.61 | 20.63 | 20.46 | 21.15 |
| Q1, Q3 | 19.42, 26.60 | 18.48, 22.80 | 19.44, 24.35 | 19.19, 25.72 |
| HIV status | | | | |
| Negative | 180 (100.0%) | 27 (100.0%) | 6 (100.0%) | 231 (100.0%) |

**Community Screening Cohort**

| | No TB (N=227) | Asymptomatic TB (N=46) | Symptomatic TB (N=7) | Total (N=290) |
|---|---|---|---|---|
| Age | | | | |
| Median | 38.69 | 34.96 | 46.95 | 37.82 |
| Q1, Q3 | 26.07, 53.67 | 25.79, 50.14 | 33.44, 52.78 | 26.28, 53.24 |
| Sex | | | | |
| Female | 113 (49.8%) | 20 (43.5%) | 4 (57.1%) | 145 (50.0%) |
| Male | 101 (44.5%) | 17 (37.0%) | 3 (42.9%) | 122 (42.1%) |
| Unknown | 13 (5.7%) | 9 (19.6%) | 0 (0.0%) | 23 (7.9%) |
| HIV status | | | | |
| Negative | 179 (78.9%) | 31 (67.4%) | 5 (71.4%) | 218 (75.2%) |
| Positive | 35 (15.4%) | 6 (13.0%) | 2 (28.6%) | 49 (16.9%) |
| Unknown | 13 (5.7%) | 9 (19.6%) | 0 (0.0%) | 23 (7.9%) |

**Clinic Triage Cohort**

| | No TB (N=212) | Symptomatic TB (N=190) | Total (N=402) |
|---|---|---|---|
| Age | | | |
| Median | 35.75 | 35.20 | 35.45 |
| Q1, Q3 | 29.10, 44.40 | 28.10, 42.25 | 28.72, 43.92 |
| Sex | | | |
| | 136 (64.2%) | 124 (65.3%) | 260 (64.7%) |

| | | | |
|---|---|---|---|
| Female<br>Male | 76 (35.8%) | 66 (34.7%) | 142 (35.3%) |
| BMI | | | |
| Median | 23.60 | 19.65 | 21.60 |
| Q1, Q3 | 20.30, 27.20 | 17.60, 23.20 | 18.70, 25.65 |
| HIV status | | | |
| Negative | 121 (57.1%) | 103 (54.2%) | 224 (55.7%) |
| Positive | 85 (40.1%) | 68 (35.8%) | 153 (38.1%) |
| Unknown | 6 (2.8%) | 19 (10.0%) | 25 (6.2%) |

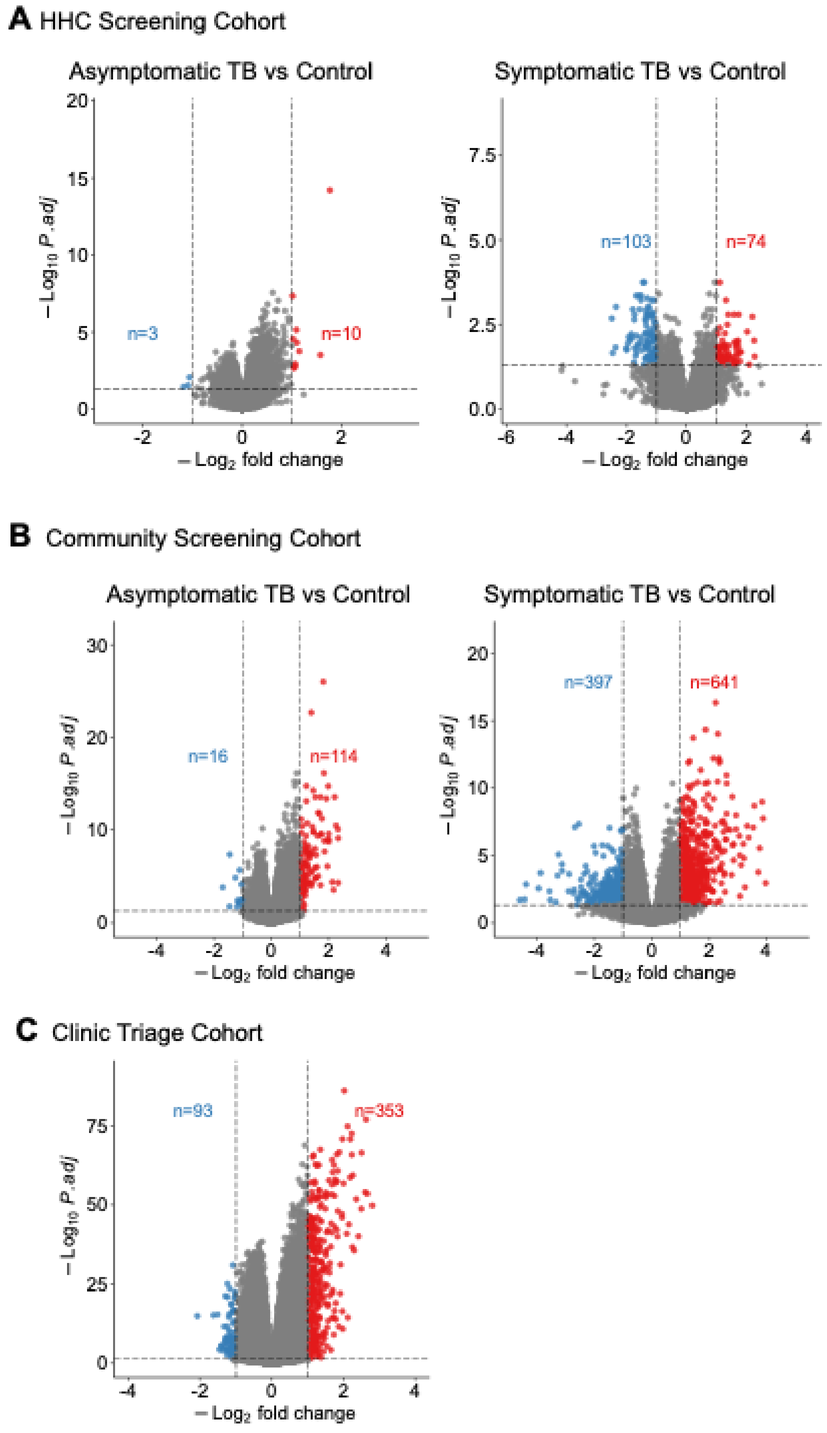


**Supplementary Figure S2. Analysis of differentially expressed genes between asymptomatic TB or symptomatic TB and TB-negative controls.** Volcano plots show the distributions of differentially expressed genes between asymptomatic TB and controls (no TB), or between symptomatic TB and controls for the Community HHC Screening (**A**), Community Screening (**B**), and Clinic Triage (**C**) cohorts. The X-axis represents $log_2$ fold change, and the Y-axis represents $-log_{10}$ of the FDR-adjusted $p$-value. Genes with significant upregulation ($log_2$ fold change ≥ 1.0, FDR p < 0.05) are highlighted in red, significant downregulation in blue, and non-significant genes in grey.

Supplementary Figure 3

**A** M3

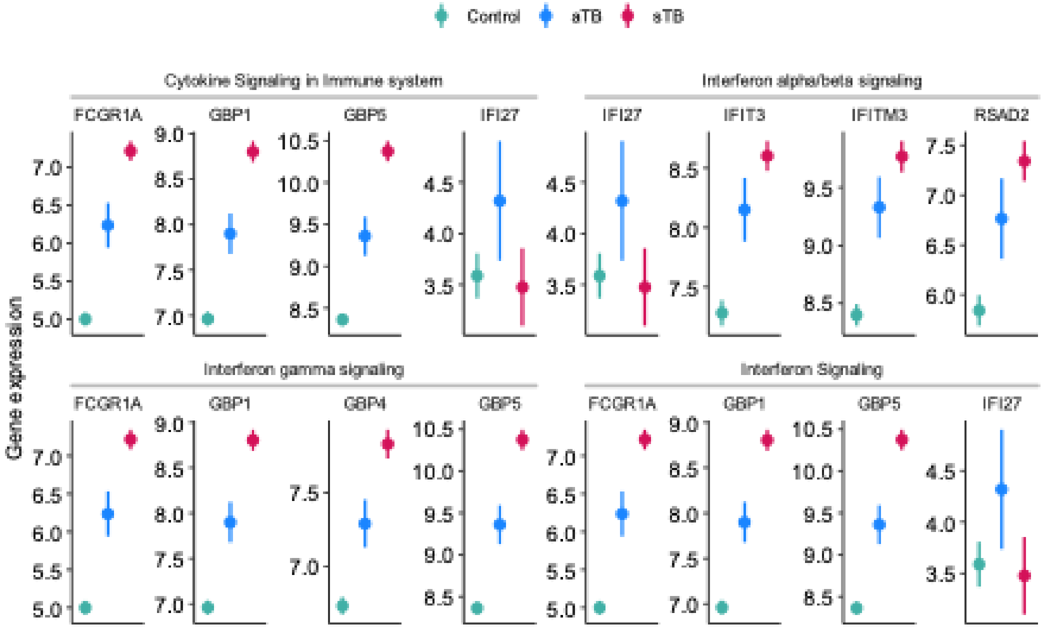


**B** M5

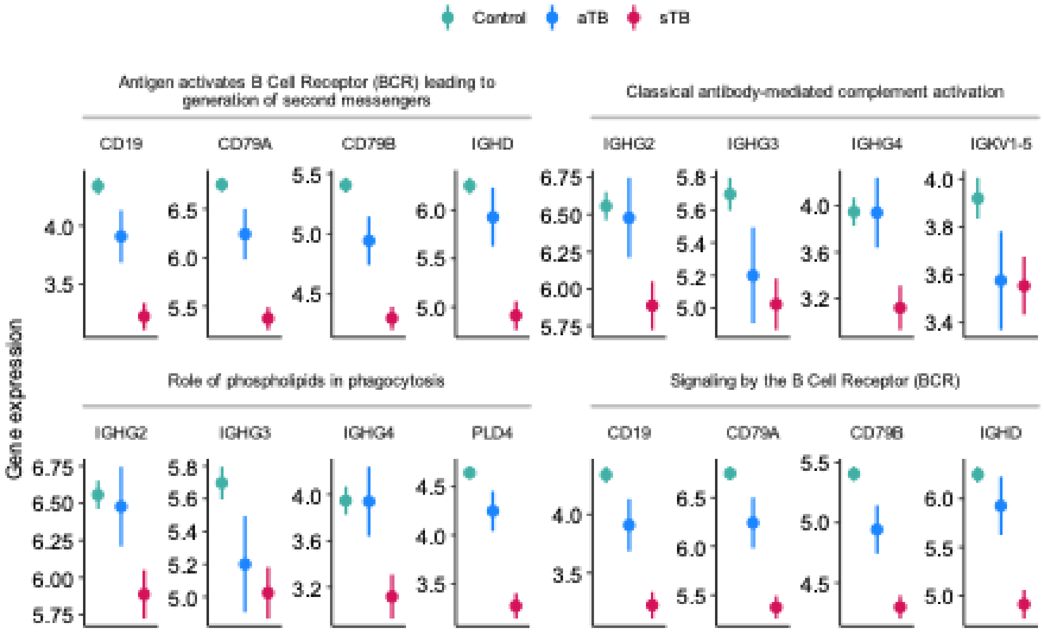

Supplementary Figure 3 continued

**C** M4

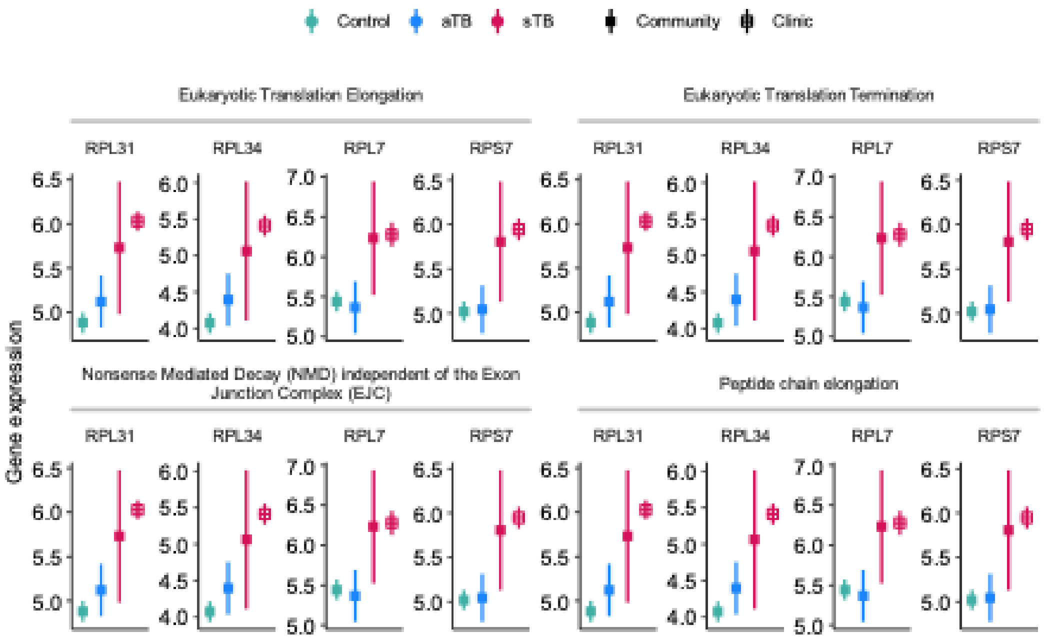


**D** M6

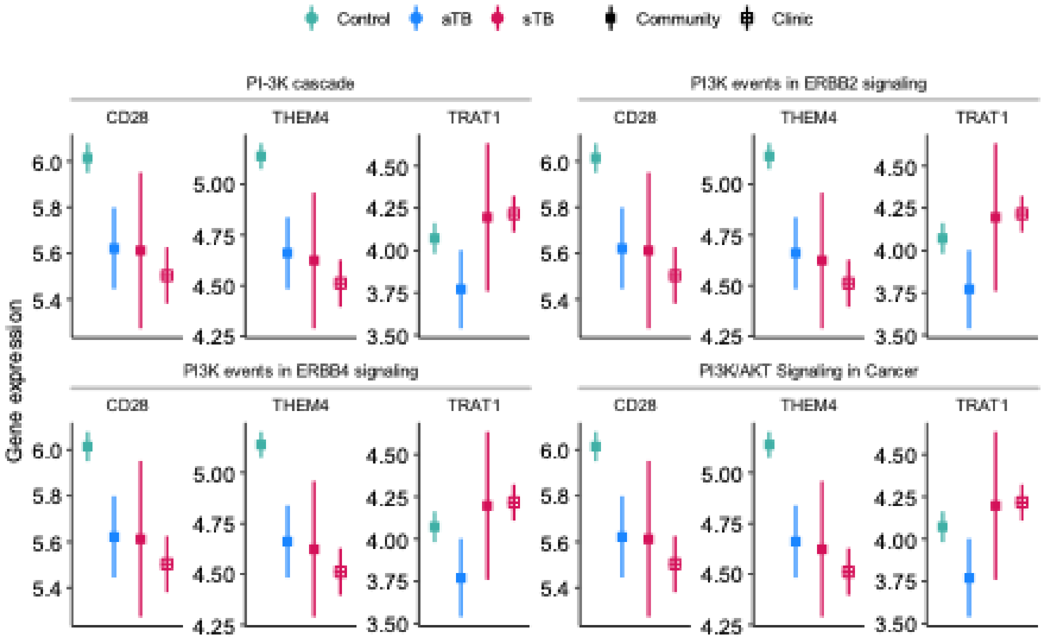

Supplementary Figure 3 continued

**E** M

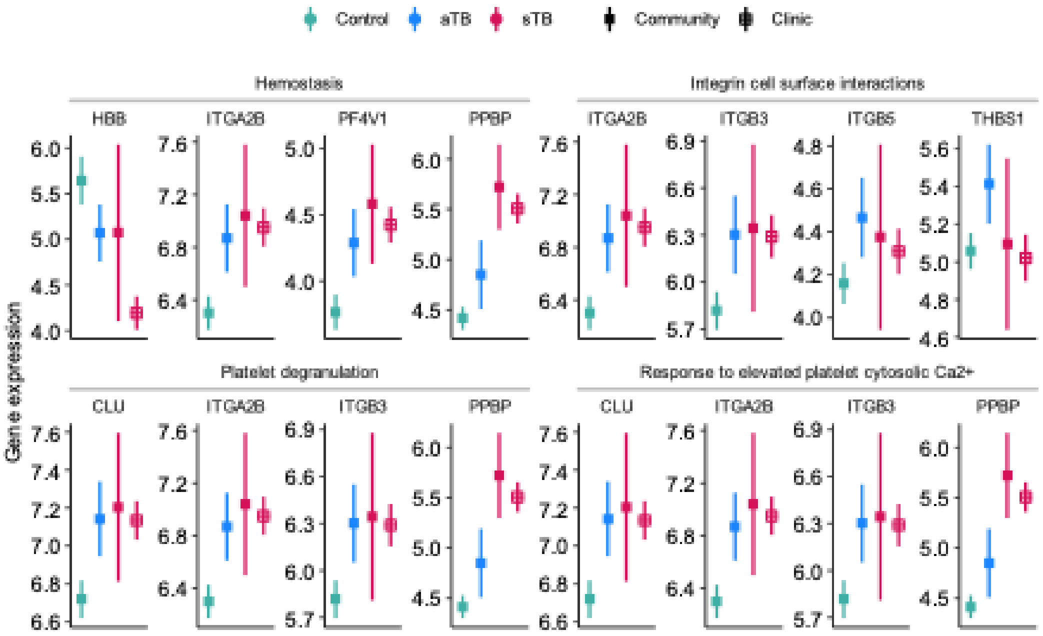


**Supplementary Figure S3. Analysis of relative expression levels for genes identified by gene co-expression network analysis across the TB spectrum.** Participants from the HHC and Community Screening and Clinic Triage cohorts were stratified into groups along the TB spectrum, from healthy controls, asymptomatic TB (aTB) and symptomatic TB (sTB, further split into Screening and Clinic Triage), by weighted gene co-expression network analysis using CEMiTool. For these analyses, data from the HHC Screening (RePORT SA) and Community Screening (CORTIS) cohorts were combined. Gene set overrepresentation analysis was performed for genes in the 6 modules identified by weighted gene co-expression network analysis. Shown are the expression levels for the top four most regulated genes from each of the top 5 significantly enriched pathways in module M3 (**A**), module M5 (**B**), module M4 (**C**), module M6 (**D**) and module M1 (**E**).

Supplementary Figure 4

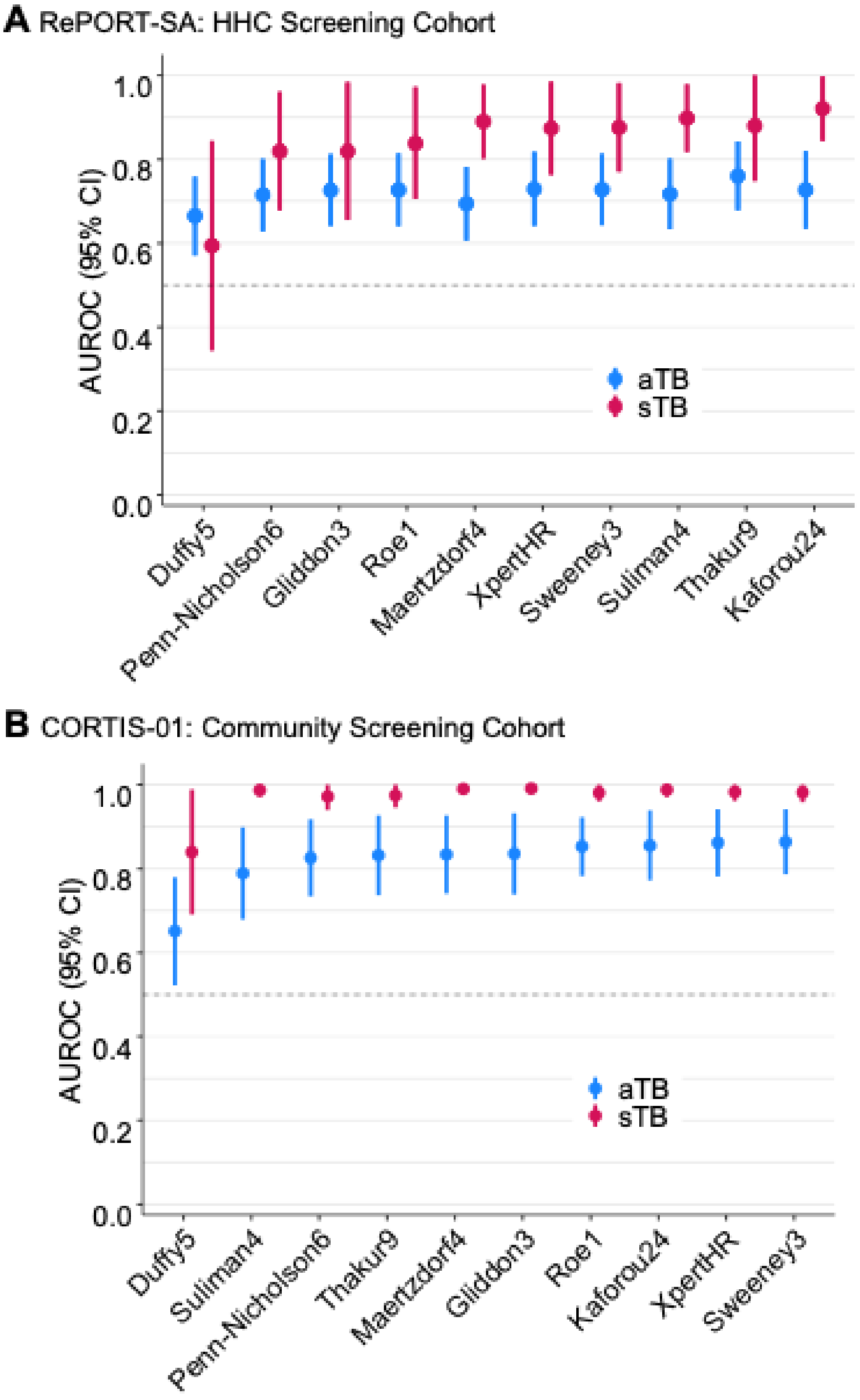


**Supplementary Figure S4. Evaluating diagnostic performance of existing, published transcriptomic biomarkers.** Area under the receiver-operating characteristic curve (AUROC) showing the performance of selected published transcriptomic signatures for differentiating between asymptomatic TB (aTB) and TB-negative controls, and between symptomatic TB (sTB) and TB-negative controls in the HHC Screening Cohort (**A**), and in the Community Screening Cohort (**B**). Shown are AUROC values (dots) and 95% CIs (error bars) for each signature.

Supplementary Figure 5

**A** HHC Screening Cohort

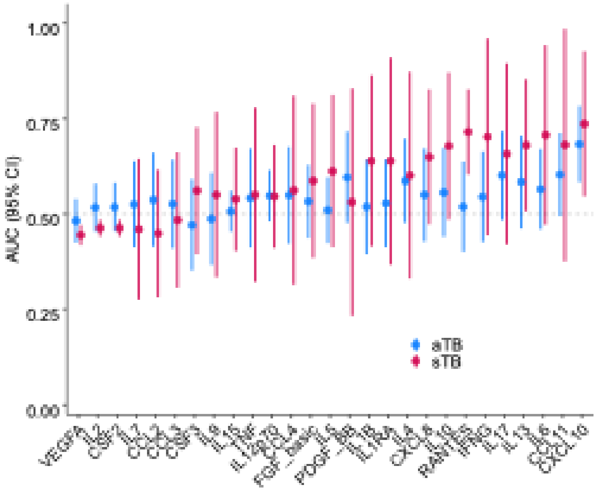


**B** Community Screening Cohort

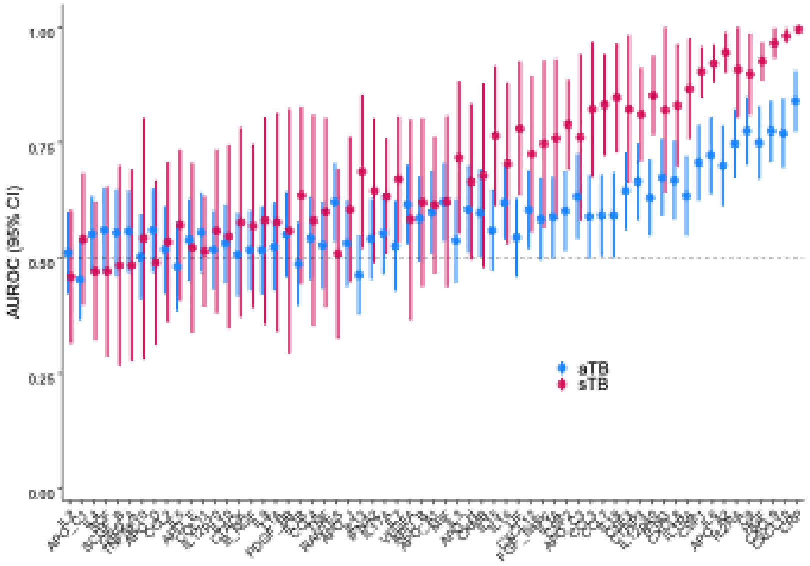


**C** Clinic Triage Cohort

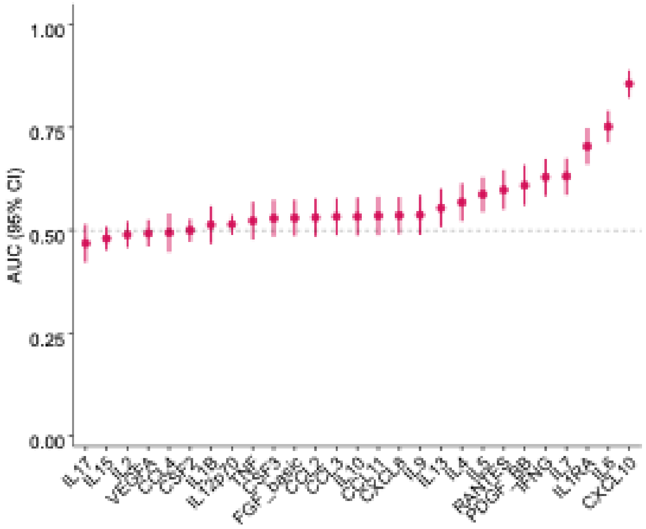


**Supplementary Figure S5. Evaluating diagnostic performance of serum proteomic markers.**

Area under the receiver-operating characteristic curve (AUROC) showing the performance of serum proteins measured by multiplex bead array in the cohorts. Diagnostic performance for differentiating between asymptomatic TB and TB-negative controls (blue), and between symptomatic TB and TB-negative controls (red) in the HHC Screening Cohort (**A**), between asymptomatic TB and TB-negative controls, and between symptomatic TB and TB-negative controls in the Community Screening Cohort (**B**), and between symptomatic TB and symptomatic TB-negative controls in the Clinic Triage cohort (**C**). Dots and error bars indicate the AUROC and 95% CIs.

Supplementary Figure 6

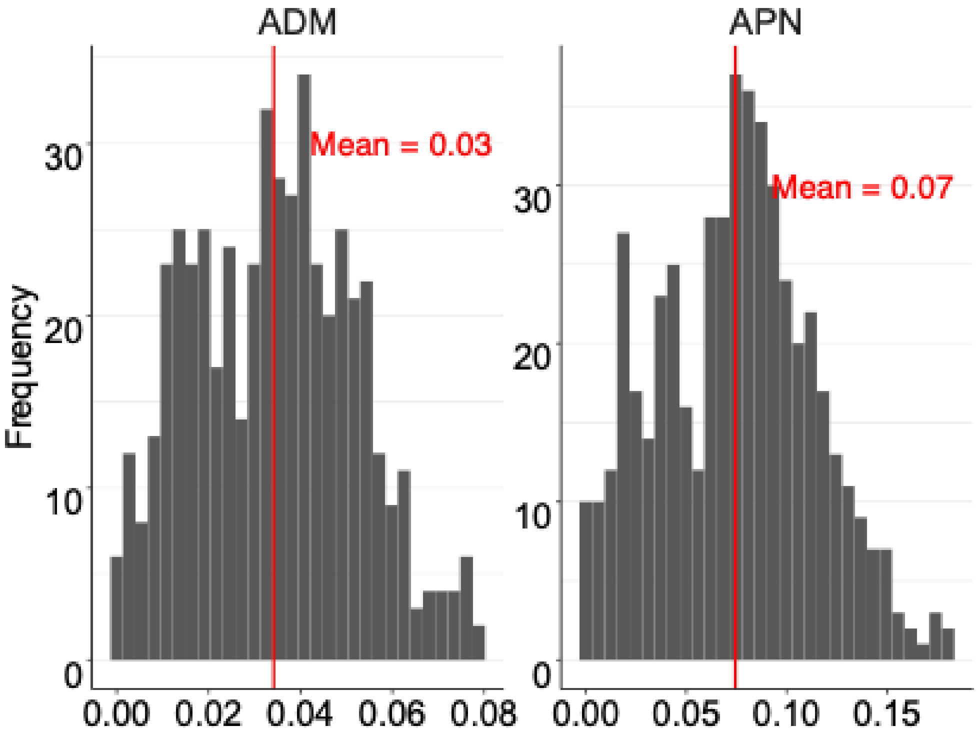


**Supplementary Figure S6. Stability analysis of asymptomatic TB clusters identified by fused similarity network analysis.** Two metrics were used to evaluate robustness of the two identified clusters among asymptomatic TB participants, estimated from 500 resampling iterations, each involving a random selection of 80% of the participant samples. (Left) Average Distance between Means (ADM), with a mean value of 0.03, indicates a small separation between resampled cluster centroids. (Right) Average Proportion of Non-overlap (APN), with a mean value of 0.07, reflects high cluster stability across resampling. Together, these low ADM and APN values suggest that the two subclusters identified by similarity network fusion are stable and reproducible.

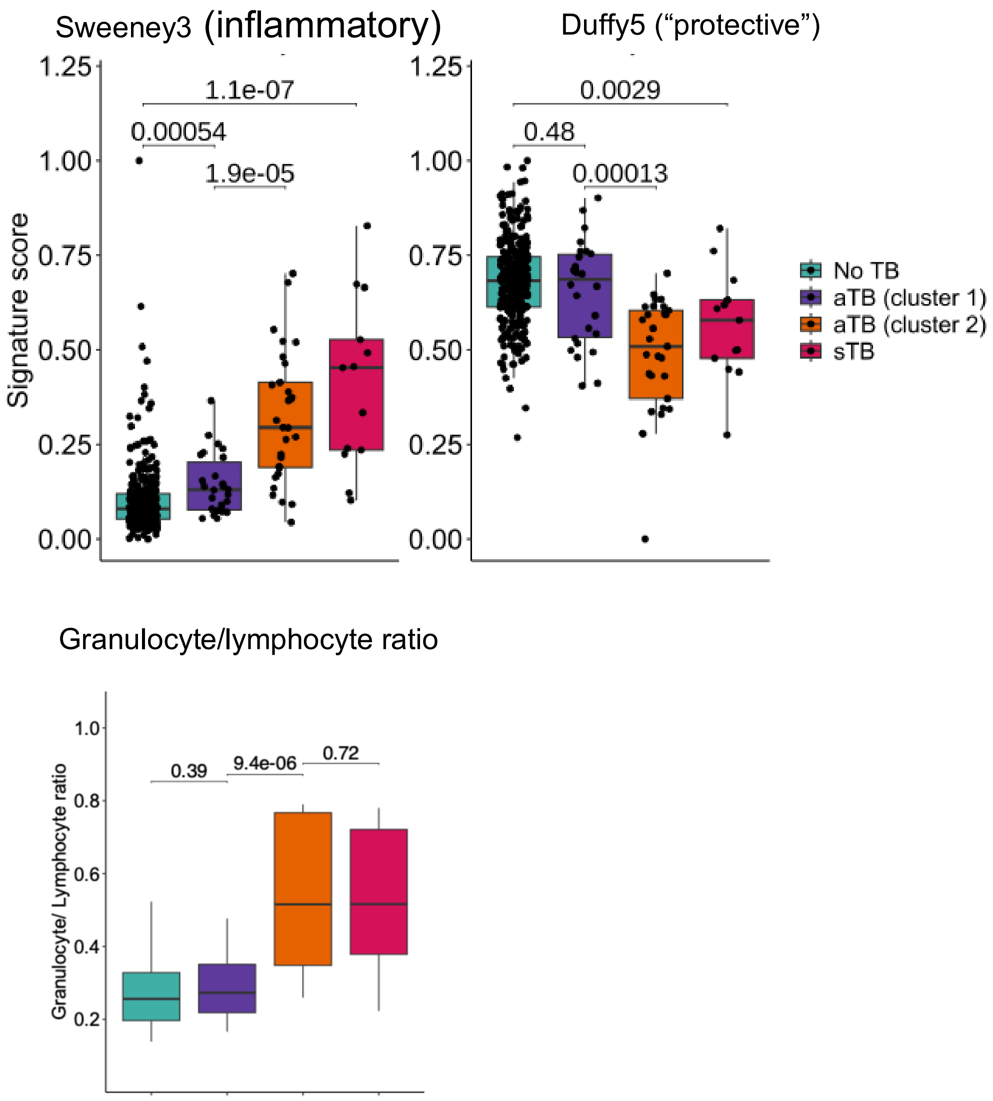


**Supplementary Figure S7. Transcriptomic signature scores for Sweeney3, Duffy5 and Granulocyte/lymphocyte ratio in cluster 1 and 2 individuals as well as No-TB controls or symptomatic TB.** The shown $p$-values were computed with the Mann-Whitney U test. For box-and-whisker plots horizontal lines indicate the median, the boxes the interquartile range (IQR) and the whiskers the range of data with $1.5 \times IQR$ from the lower and upper quartiles.

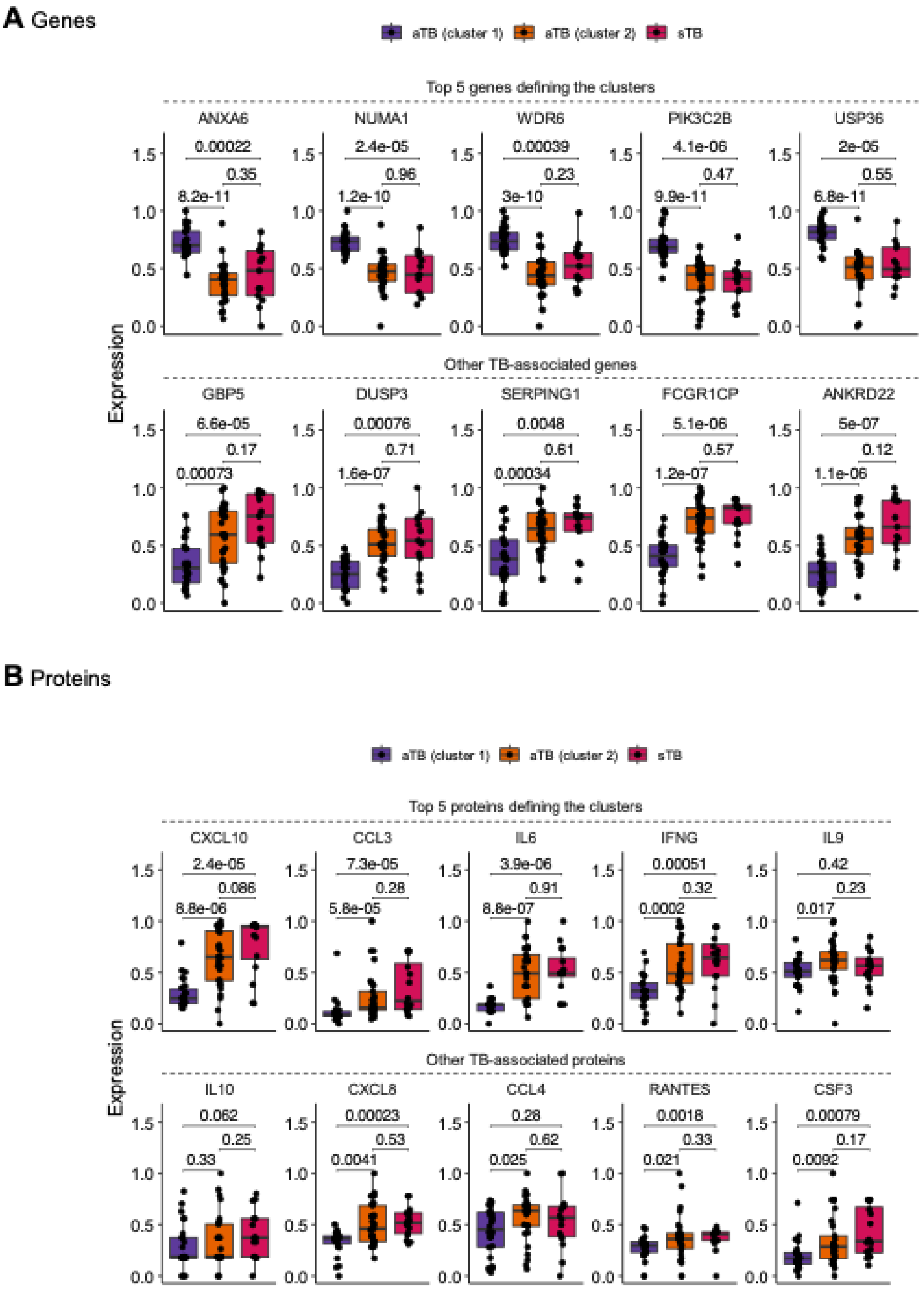


**Supplementary Figure S8. Analysis of expression levels of the top genes and proteins defining the two asymptomatic TB clusters obtained from fusion similarity network analysis.** (**A**) Expression levels of the top 5 genes defining the subclusters (top), contrasted with the expression levels of selected 5 known TB-associated genes. (**B**) Expression levels of the top 5 proteins defining the subclusters of asymptomatic TB (aTB) (top), contrasted with the expression levels of other selected 5 proteins measured in the samples.
